# Zinc and iron homeostatic interactions in a mutant lacking nicotianamine vacuolar storage and citrate xylem loading

**DOI:** 10.64898/2026.03.19.712977

**Authors:** Steven Fanara, Maxime Scheepers, Madeleine Boulanger, Marie Schloesser, Bernard Bosman, Monique Carnol, Anthony Fratamico, Manon Sarthou, Pierre Tocquin, Marc Hanikenne

**Author notes:** Plant Science Research Laboratory (LRSV), UMR5546 CNRS/University of Toulouse/Toulouse-INP, 24 Chemin de Borde Rouge, 31320 Auzeville Tolosane, France. GDTech S.A. avenue de l’Expansion, 7, 4432 Alleur, Liège, Belgium. These two authors contributed equally to this work (Steven Fanara, Maxime Scheepers). **Contact** Steven Fanara and Marc Hanikenne.

## Abstract

Metal homeostasis in plants relies on coordinated uptake, chelation, and transport mechanisms involving, but not limited to, citrate and nicotianamine (NA). In Arabidopsis (*Arabidopsis thaliana*), disruption of the citrate exporter FRD3 (FERRIC REDUCTASE DEFECTIVE 3) causes constitutive Fe deficiency responses, altered iron (Fe), manganese (Mn) and zinc (Zn) distribution, with Fe accumulation in the root cell wall. This ultimately results in oxidative and biotic stress responses, and impaired root development, phenotypes that are partially alleviated by Zn excess. In this study, we investigated the consequences of impairing both citrate loading into xylem vessels and NA partitioning within cells. The *frd3 zif1* double mutant exhibits enhanced sensitivity to Zn excess, severe defects in root system architecture and meristem maintenance, persistent oxidative stress, and compromised reproductive development. These phenotypes correlate with sustained activation of Fe deficiency signaling and marked defects in root-to-shoot metal translocation. Our findings reveal that coordinated citrate export and NA compartmentation form an integrated buffering strategy required to maintain metal homeostasis and partitioning, as well as redox balance and proper development, including root plasticity and seed yield, under fluctuating metal availability.

## Introduction

Living organisms require optimal concentrations of a number of essential nutrients to complete their life cycle. Among these essential nutrients are found metal ions such as iron (Fe), manganese (Mn), and zinc (Zn). Metals are required for numerous biological functions. In plants, metals play for instance critical roles in maintaining protein structure, as co-factors of enzymatic activities, or as key components of the photosynthetic and respiratory processes. Metals are needed in small quantities for correct growth and development but become toxic when accumulating in tissues in large amounts. Plants have developed tightly regulated homeostatic mechanisms to maintain optimal metal concentrations in tissues in a range of habitats with varying metal supply (Hänsch and Mendel, 2009; Palmer and Guerinot, 2009).

Although Fe is one of the most abundant metals in the Earth’s crust, it is hardly accessible from soil for plants especially at alkaline pH (Schmidt et al., 2014; Rengel et al., 2023). Dicotyledonous plants, such as *Arabidopsis thaliana*, have developed a Fe reduction strategy (reduction of Fe^3+^ to Fe^2+^, also called strategy I) to efficiently take up Fe from the soil (Schmidt, 2003; Walker and Connolly, 2008; Hanikenne et al., 2021; Liang, 2022). In *A. thaliana*, the first step of strategy I is the acidification of the rhizosphere to increase Fe solubility, which is enabled by the ARABIDOPSIS H^+^-ATPase 2 (AHA2) that is expressed in the plasma membrane of root epidermal cells (Guerinot and Yi, 1994; Robinson et al., 1999; Santi and Schmidt, 2009). Fe solubilization is further increased by the secretion of phenolic compounds such as coumarins (Fourcroy et al., 2014; Schmidt et al., 2014; Robe et al., 2021; Robe et al., 2025). Fe^3+^ is then reduced to Fe^2+^ by FERRIC REDUCTION OXIDASE 2 (FRO2) (Robinson et al., 1999). Finally, Fe^2+^ is transported into roots by IRON-REGULATED TRANSPORTER 1 (IRT1), which is the main Fe transporter (Eide et al., 1996; Vert et al., 2002). At the transcriptional level, *FRO2* and *IRT1* genes are under the control of BASIC HELIX-LOOP-HELIX PROTEIN (bHLH) transcription factors, including FIT (FER-LIKE IRON DEFICIENCY-INDUCED TRANSCRIPTION FACTOR; bHLH29) and other associated bHLH proteins that cooperatively up-regulate their expression upon Fe deficiency (Colangelo and Guerinot, 2004; Yuan et al., 2005; Gratz et al., 2021; Spielmann et al., 2023).

IRT1 has a high affinity for Fe^2+^ but has a low specificity as it transports other divalent metal cations such as Mn^2+^, Zn^2+^ or Cd^2+^ (Eide et al., 1996; Vert et al., 2002; Spielmann et al., 2022; Cointry et al., 2025). Therefore, these metals are also taken up in large amounts upon Fe deficiency, and a number of genes involved in the vacuolar storage of Mn, nickel (Ni) or Zn are also upregulated in a FIT-dependent manner (Arrivault et al., 2006; Haydon et al., 2012; Eroglu et al., 2016; Schwarz and Bauer, 2020). Other genes (e.g., *ZINC INDUCED FACILITATOR 1*, *ZIF1*) are controlled by additional regulators of Fe homeostasis such as POPEYE (PYE), or MYB10 and MYB72 (MYELOBLASTOSIS) (Long et al., 2010; Palmer et al., 2013; Wu et al., 2024).

Numerous pieces of evidence point to an interconnection between Fe and Zn homeostasis (Haydon and Cobbett, 2007b; Shanmugam et al., 2011; Haydon et al., 2012; Charlier et al., 2015). Plants exposed to Zn excess have a phenotype similar to plants exposed to Fe deficiency: shoot chlorosis, decreased growth, limited Fe root concentration, but without a significant decrease of shoot Fe concentration (Fukao et al., 2011; Lešková et al., 2017; Rengel et al., 2023; Thiébaut et al., 2025b). Zn excess toxicity can be suppressed by an increased Fe supply (Shanmugam et al., 2011). Hence, Zn excess mimics Fe deficiency at both phenotypic and transcriptomic levels as many Fe homeostasis actors are overexpressed upon Zn excess, including not only *FIT* and *MYB72* (Lešková et al., 2017), but also *FRO2* and *IRT1* (Thiébaut et al., 2025b).

Several genes having a role in the Zn and Fe interaction have been identified. One of these genes is *ZIF1*, a vacuolar transporter of nicotianamine (NA) involved in Fe and Zn homeostasis (Haydon and Cobbett, 2007b). *ZIF1* is up-regulated upon Zn excess or Fe deficiency and has been proposed to detoxify Zn by enabling its vacuolar storage in these conditions (Haydon et al., 2012).

The citrate plasma membrane transporter FERRIC REDUCTASE DEFECTIVE 3 (FRD3) is another important actor of the Zn and Fe homeostasis interaction. FRD3 is expressed in root pericycle cells and is involved in citrate loading into xylem vessels. Citrate is necessary for a correct distribution of Fe in roots, shoots, anthers, pollen and seeds (Rogers and Guerinot, 2002; Green and Rogers, 2004; Durrett et al., 2007; Roschzttardtz et al., 2011). *FRD3* is also involved in Zn tolerance (Pineau et al., 2012), and displays a complex regulation in response to Zn in *A. thaliana* (Charlier et al., 2015). A *frd3* mutant displays a strongly impaired Fe homeostasis, including Fe mis-localization in tissues and constitutive Fe deficiency response (Rogers and Guerinot, 2002; Green and Rogers, 2004; Durrett et al., 2007; Roschzttardtz et al., 2011). Our recent study suggested that this altered Fe distribution triggers cell wall (CW) modifications linked to oxidative and biotic stress responses (Scheepers et al., 2020). It also revealed that Mn toxicity has a major contribution to the mutant phenotype. Finally, it was observed that Zn excess inhibited both oxidative and biotic responses and partially restored root growth of the mutant as well as citrate and NA levels in roots (Scheepers et al., 2020).

Citrate and NA are chelating compounds that form soluble complexes with Fe, Mn and Zn and that facilitate their transport at both the subcellular and whole-plant level (Curie et al., 2009; Morrissey and Guerinot, 2009; Flis et al., 2016; Seregin and Kozhevnikova, 2023; Hollmann et al., 2025). Additionally, Fe and Zn ligands can be almost instantaneously exchanged between citrate and NA (Rellán-Álvarez et al., 2008; Rellán-Álvarez et al., 2010). Levels of citrate are increased in the xylem sap of a biosynthetically NA-deficient mutant (*nas4x*, lacking the four nicotianamine synthase proteins) (Schuler et al., 2012) Consistently, the expression of *FRD3* is highly induced in *nas4x* mutant, whereas *NAS* genes are overexpressed in *frd3* mutant roots (Schuler et al., 2012). In particular, the *frd3* mutant displays *NAS2* gene over-expression in control condition but not upon Zn excess (Scheepers et al., 2020). Moreover, *nas4x* mutant plants display strong Fe deficiency symptoms, including the up-regulation of the Fe uptake machinery genes (Klatte et al., 2009; Schuler et al., 2012), similarly to *frd3* mutant (Scheepers et al., 2020) or *ZIF1* overexpressors (Haydon et al., 2012). Indeed, *ZIF1* overexpression lines exhibit increased vacuolar sequestration of NA in roots, leading to NA depletion in leaves. This is accompanied by a vacuolar Zn but not Fe sequestration in roots and by a reduced lateral mobility of Fe in the leaf blade, ultimately triggering constitutive Fe deficiency symptoms (Haydon et al., 2012). Interestingly, *frd3 nas4x* mutants display a more severe interveinal leaf chlorosis (which aggravates with age) compared to *frd3* or *nas4x* mutants (Schuler et al., 2012). The study of the *frd3 nas4x* quintuple mutant depicted a role for citrate in xylem-fed tissues and for NA in Fe remobilization out of the phloem, and therefore in phloem-fed tissues (Schuler et al., 2012). Finally, in a *frd3* mutant, *ZIF1* is overexpressed, which might contribute to the intracellular relocation of both Fe and Zn that are excessively accumulating in *frd3* roots (Scheepers et al., 2020).

In this study, we investigated the interplay between xylem sap loading of citrate and translocation of metal-citrate complexes to shoots (achieved through FRD3 activity) and metal excess detoxification achieved by metal-NA sequestration into root vacuoles through ZIF1 activity. To address the question of how increased cytosolic NA levels (*zif1* mutation) influence the mobility of Zn and other NA-chelated metals (e.g., Fe) in the context of impaired citrate-dependent xylem loading (*frd3* mutation), we generated *frd3 zif1* double mutants as a genetic tool. By integrating molecular, ionomic, biochemical and developmental analyses across vegetative and reproductive stages, we sought to comprehensively assess the impact of disrupted chelator compartmentation on metal homeostasis.

## Results

### frd3 zif1 double mutants present distinct root system architecture, particularly upon Zn excess

To unravel how elevated cytosolic nicotianamine (NA) levels influence the mobility of Zn and other NA-chelated metals, such as Fe, in an impaired citrate xylem loading context, *frd3-7 zif1* double mutants were generated. Two *zif1* T-DNA insertion lines were selected and crossed with *frd3-7* (Scheepers et al., 2020): *zif1-2* (Haydon and Cobbett, 2007a), with a T-DNA insertion at the 5’UTR-exon 1 junction, and a so far unexamined allele, *zif1-6*, with a T-DNA insertion in the intron 9-exon 9 junction region **(Supplementary Figure S1A-B)**. Prior crossing with *frd3-7*, the Zn sensitivity phenotypes of these two *zif1* mutants were assessed in our experimental conditions. Consistent with previous reports (Haydon and Cobbett, 2007a; Lee et al., 2021), both *zif1* mutants were significantly more affected than the WT (Col-0) under Zn excess conditions **(Supplementary Figure S1C-F)**, whereas *zif1* single mutants displayed similar root length **(Supplementary Figure S1C-D)**, shoot fresh weight per plant **(Supplementary Figure S1E)** and total chlorophyll content per plant **(Supplementary Figure S1F)** than the WT in control conditions. Altogether, this indicates that the *zif1-6* allele is phenotypically similar to the previously described *zif1-2* allele (Haydon and Cobbett, 2007a).

To investigate how Zn availability affects root development, we performed a detailed analysis of root system architecture under Zn excess (**Figure 1A**). Under control conditions, primary root (PR) growth in *frd3-7* and *frd3-7 zif1* mutant plantlets was altered, with the double mutants displaying the shortest primary roots, whereas *zif1* single mutants did not show any growth defect (**Figure 1B**). High Zn conditions (75 µM Zn) negatively affected root growth of all genotypes, resulting in significantly shorter roots in all single and double mutants compared with the WT (**Figure 1B**). In particular, *frd3-7 zif1* double mutants were the most severely affected, with a ∼67% PR length reduction compared to the control condition, whereas *frd3-7* and *zif1* mutant roots were reduced by approximately 10.5% and 33%, respectively, reaching comparable lengths (**Figure 1B and Supplementary Table S1)**.

**Figure 1.**
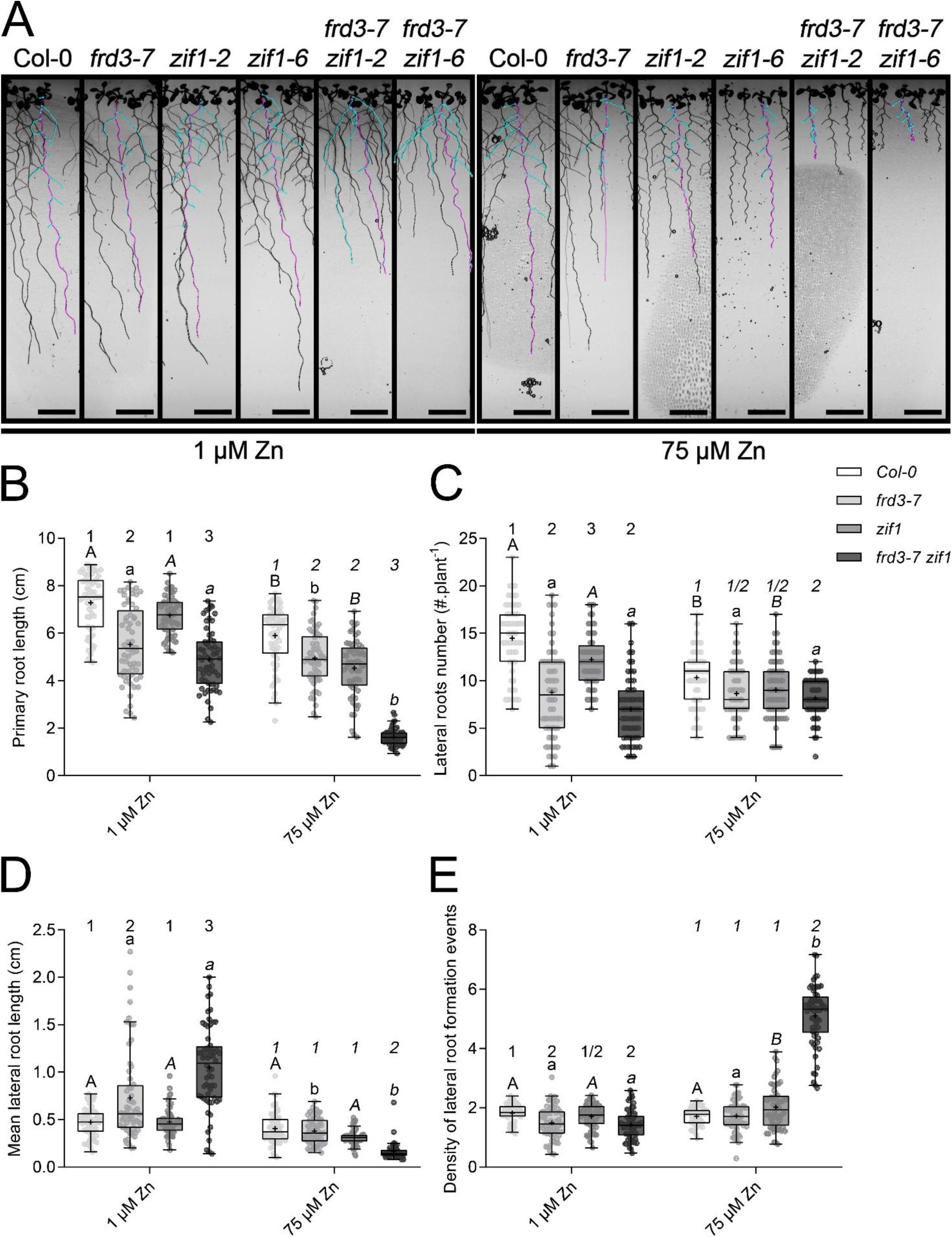
Root system architecture upon zinc excess in *frd3-7 zif1* double mutants. **(A)** Traces of primary roots (magenta) and lateral roots (cyan) of a representative individual of Col-0, *frd3-7*, *zif1* and *frd3-7 zif1* plantlets after 14 days of growth on solid media containing different Zn concentrations (1 and 75 µM ZnSO_4_). Scale bars: 1 cm. **(B)** Primary root length, **(C)** lateral root number per primary root shown in **(B)**, **(D)** mean lateral root length for each primary root shown in **(B)**, and **(E)** density of lateral root formation events [defined as lateral root number per cm of primary root shown in **(B)**] of Col-0 (white), *frd3-7* (light grey), *zif1* (middle grey) and *frd3-7 zif1* (dark grey) plantlets after 14 days of growth on solid media containing different Zn concentrations (1 and 75 µM ZnSO_4_). Box-and-whisker plots show the median (line), interquartile (box), 1.5 interquartile (whiskers), and outliers (points beyond the whiskers) of values from 4 independent experiments each including 3 series of 8 plantlets per genotype and condition. Mean values are represented by ‘+’ symbols. The data were analyzed by two-way ANOVA and followed by Bonferroni multiple comparison post-test. Statistically significant differences between conditions are indicated by letters (*P*<0.05) or between genotypes are indicated by digits (*P*<0.05). Images were acquired using a Petrimaton apparatus. All features were measured using the *simple neurite tracer* and *cell counter* plugins on ImageJ/FIJI, respectively. All mentions of lateral roots refer to 1^st^ order lateral roots only. All mean, median and relative values are provided in **Supplementary Table S1**.

Expanding the characterization of the root system architecture to lateral roots (LR) led to interesting observations. First, *frd3-7*, *zif1* and *frd3-7 zif1* already showed lower number of LR in control conditions (**Figure 1C**), although LR lengths were longer in *frd3-7* and *frd3-7 zif1* (**Figure 1D**). This lower number of LR resulted in lower densities of LR (number per cm of PR) in *frd3-7* and *frd3-7 zif1* in control conditions (**Figure 1E**). Second, exposure to Zn excess markedly decreased (i) the number of LR in the WT (**Figure 1C**) and (ii) the mean length of LR in *frd3-7* (**Figure 1D**), alleviating differences between WT and the *frd3-7* and *zif1* single mutants. In contrast, *frd3-7 zif1* double mutants exhibited the shortest LR under Zn excess, while LR density was strongly increased (**Figure 1E**) due to the severe inhibition of the PR growth (**Figure 1B**) combined with the maintenance of a similar LR number (**Figure 1C**).

Together, these results indicate that primary and lateral root development are strongly modulated by Zn availability and metal homeostasis status. While Zn excess led to more pronounced and persistent modifications of root system architecture in a double mutant impaired in several metal chelator pathways (as in the *frd3-7 zif1*), it partially alleviated lateral root alterations in the *frd3-7* single mutant. This highlights a differential sensitivity of root developmental responses to metal stress across genotypes that are distinctively defective in metal chelators partitioning.

### Zn excess exacerbates shoot growth defects, chlorosis, and ionomic alterations in frd3 zif1 double mutants

Under control conditions, the shoot fresh weight was already reduced in *frd3-7 zif1* mutants compared with the WT, whereas *frd3-7* single mutant showed only a mild decrease **(Supplementary Figure S2A)**. Exposure to Zn excess significantly decreased shoot fresh weight in all mutants, with the *frd3-7 zif1* double mutants exhibiting the most pronounced reduction **(Supplementary Figure S2A)**. Total chlorophyll content was lower in all mutants relative to the WT in control conditions, with *frd3-7 zif1* double mutants displaying the most severe chlorosis (**Supplementary Figure S2B**). Under Zn excess, chlorosis symptoms were comparable between the WT and *zif1* single mutants, and *frd3-7* and *frd3-7 zif1* displayed the strongest reduction in chlorophyll content **(Supplementary Figure S2B)**.

To better understand whether the aggravated growth defects observed in *frd3-7 zif1* double mutants were associated with altered metal homeostasis, we next quantified Fe, Mn, and Zn concentrations in whole plantlets **(Supplementary Figure S3A-C)**. Under control conditions, the reduced growth of *frd3-7 zif1* double mutants was accompanied by a significant accumulation of Fe and Mn, but not Zn, compared to the WT and single mutants. Under Zn excess, *frd3-7 zif1* plantlets maintained elevated Fe and Mn levels **(Supplementary Figure S3A-B)**, whereas Zn concentrations remained comparable to those of the WT and the *frd3-7* single mutant **(Supplementary Figure S3C)**. In contrast, *frd3-7* and *zif1* single mutants displayed metal concentrations similar to the WT under both conditions **(Supplementary Figure S3A-C)**.

### Prolonged and severe Zn excess impacts root apical meristem (RAM) development

Given the strong modifications in root system architecture observed in the double mutants, we next examined the integrity of the primary root apical meristem (RAM) under different mineral conditions, including two conditions of Zn excess (75 and 150 µM Zn). Inhibition of primary root growth, together with the maintenance of a relatively high lateral root density, suggests impairment of the primary RAM and establishment of compensatory mechanisms aiming at sustaining the uptake of essential minerals. Thus, we measured RAM size using two metrics: the length in micrometers (µm) and in number of cells in the cortex cell layer. The RAM length was assessed as the distance between the quiescent center and the first elongated cortex cell (lower and upper red arrows in **Figure 2A**, respectively) (Thiébaut et al., 2025b). Under control conditions, all genotypes presented similar RAM length, except for *frd3-7 zif1* double mutants that showed reduced RAM size (**Figure 2A-C**). Similarly to short-term Zn excess (Thiébaut et al., 2025b), exposure to Zn excess (during two weeks from germination) led to shortened RAM in all genotypes, with clear tendencies of increased inhibition of RAM size in all mutants compared to the WT. Finally, accentuating Zn excess (e.g., 150 µM Zn) led to even shorter RAM in all genotypes, with visible entry of propidium iodide within dead cells (**Figure 2A**), suggesting that cell death was induced in *zif1* single and *frd3-7 zif1* double mutants but not in the WT or *frd3-7*. These observations indicate that *zif1* and *frd3-7 zif1* mutants were hypersensitive to severe Zn excess. Notably, although RAM length (in µm) was comparable among genotypes at 150 µM Zn, the number of cortex cells was markedly reduced in *zif1* single and *frd3-7 zif1* double mutants (**Figure 2B**). This suggests that the maintenance of a similar RAM size likely results from increased cortex cell size rather than sustained cell proliferation.

**Figure 2.**
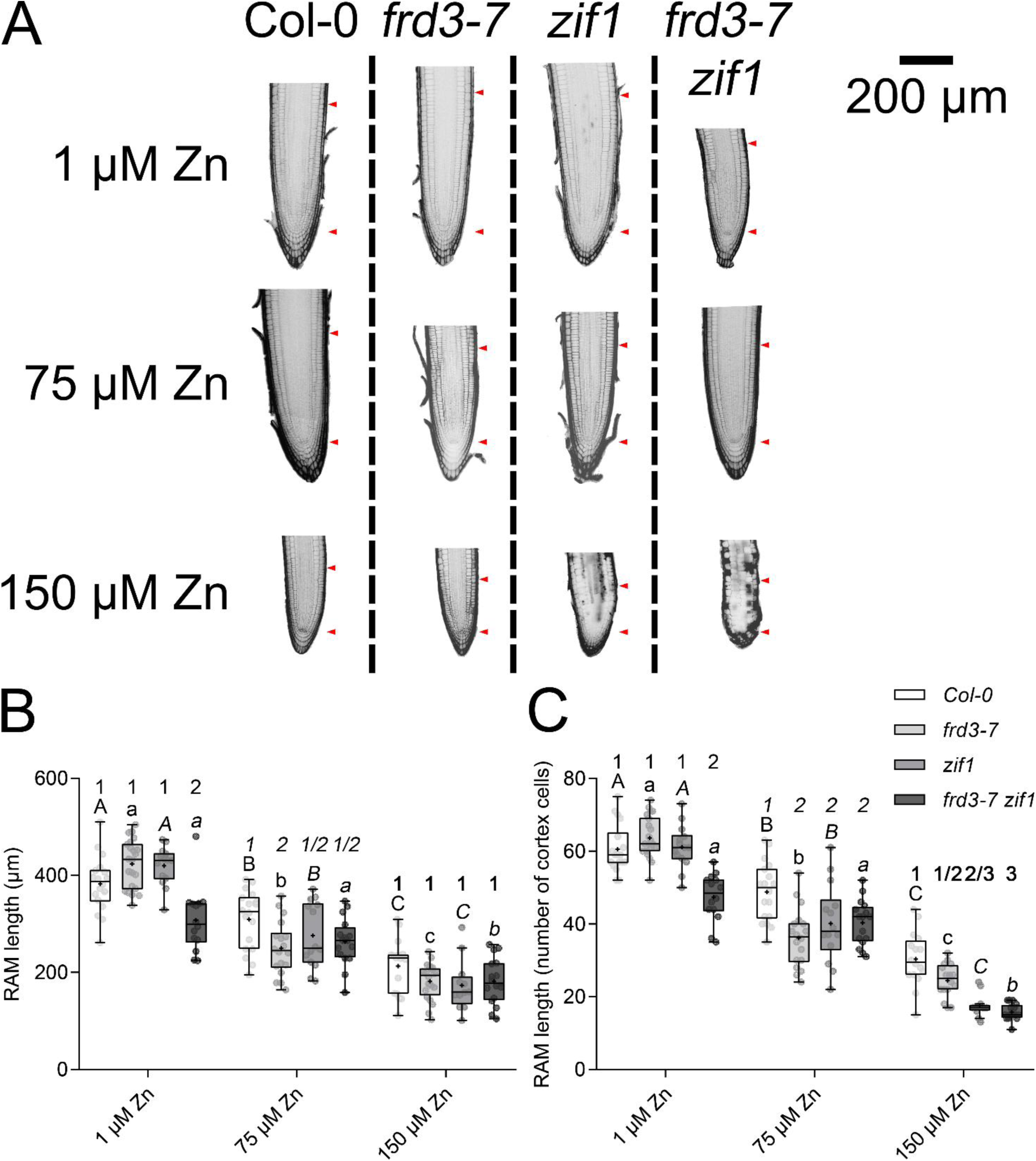
Impact on the root apical meristem (RAM) of zinc excess in *frd3-7 zif1* double mutants. **(A)** Representative images of the RAM of Col-0, *frd3-7*, *zif1* and *frd3-7 zif1* plantlets after 14 days of growth on solid media containing different Zn concentrations (1, 75 and 150 µM ZnSO_4_). RAM size was measured between the two red arrows pointing to the quiescent center (QC) (lower arrow) and the first elongated cortex cell (upper arrow), with the last non-elongated cortex cell defined as the last cell whose length was at least twofold smaller than the length of the following five cells. Images are representative of 2 independent experiments each including 5-8 plantlets per genotype and condition. **(B-C)** Length of the root apical meristem (RAM), assessed between red arrows both in micrometers (µm) **(B)** and in number of cortex cells **(C)**, of Col-0 (white), *frd3-7* (light grey), *zif1* (middle grey) and *frd3-7 zif1* (dark grey) plantlets after 14 days of growth on solid media containing different Zn concentrations (1, 75 and 150 µM ZnSO_4_). Box-and-whisker plots show the median (line), interquartile (box), 1.5 interquartile (whiskers), and outliers (points beyond the whiskers) of values from 2 independent experiments each including 5-8 plantlets per genotype and condition. Mean values are represented by ‘+’ symbols. The data were analyzed by two-way ANOVA and followed by Bonferroni multiple comparison post-test. Statistically significant differences between conditions are indicated by letters (*P*<0.05) or between genotypes are indicated by digits (*P*<0.05).

### Organic acids and nicotianamine accumulate differentially in the different genotypes

The pronounced alterations in RAM organization under Zn excess, particularly in genotypes defective in NA vacuolar sequestration, prompted us to examine whether these developmental defects were associated with changes in the intracellular balance of key metal chelators. Considering their key roles in metal cellular mobility, we next quantified citrate, malate, and nicotianamine (NA) levels in roots of the WT, *frd3-7* and *zif1* single mutants, and *frd3-7 zif1* double mutants **(Supplementary Figure S4)**. Under control conditions, *frd3-7 zif1* double mutants exhibited intermediate citrate and malate levels compared with *frd3-7* roots, which accumulated substantially higher amounts of all three chelators relative to the WT. Indeed, *frd3-7* roots contained ∼3.3-fold more citrate **(Supplementary Figure S4A)**, ∼3.1-fold more malate **(Supplementary Figure S4B)**, and ∼11.5-fold more NA **(Supplementary Figure S4C)**. By contrast, *zif1* single mutants displayed comparable levels of the three chelators to the WT **(Supplementary Figure S4A-C)**.

Under severe Zn excess, citrate levels in roots of *frd3-7 zif1* double mutants were comparable to those of the WT and *frd3-7* **(Supplementary Figure S4A)**, whereas malate levels were elevated (∼2-fold higher than WT), resembling the *zif1* single mutant **(Supplementary Figure S4B)**. In *frd3-7* roots, citrate and malate concentrations decreased to WT-like levels **(Supplementary Figure S4A-B)**. NA concentrations strongly decreased in *frd3-7* roots upon severe Zn excess, and although levels also declined in the double mutants, they remained higher than in *frd3-7*, reaching amounts similar to the WT, but lower than *zif1* single mutants **(Supplementary Figure S4C)**.

These observations show that chelator levels are differentially affected depending on the genetic background and Zn supply, with *frd3-7 zif1* double mutants displaying intermediate accumulation patterns under control conditions and a distinct profile under Zn excess.

### The Fe uptake machinery is strongly and constitutively induced in plantlet roots of frd3 zif1 double mutants

As alterations of the root system architecture observed in the double mutants are accompanied by RAM size reduction and high accumulation of Fe in plantlets, whose uptake is primarily IRT1-dependent, we next investigated the activation of the Fe uptake machinery (relying on AHA2, FRO2 and IRT1). We assessed the activity and abundance of these proteins in roots of plantlets grown under control and Zn excess conditions. Consistent with transcriptomics data obtained upon short-term Zn excess (Thiébaut et al., 2025b), the activity of the Fe uptake machinery was induced by 14 days of Zn excess, including AHA2 H^+^-ATPase activity and FRO2-driven ferric chelate reductase activity (**Figure 3A-B**). A rhizosphere acidification assay showed increased AHA2 activity in *frd3-7* and *frd3-7 zif1* under control conditions, which was further enhanced by Zn excess (**Figure 3A**). In contrast, no clear induction of AHA2 activity was detected in *zif1* single mutants (**Figure 3A**). Similarly, FRO2 activity was induced in *frd3-7* and *frd3-7 zif1* under control conditions, which was accentuated upon exposure to Zn excess (**Figure 3B**). Consistent with these observations, IRT1 protein abundance was visibly higher in *frd3-7* and *frd3-7 zif1* than in the WT and *zif1* single mutants under control conditions (**Figure 3C**), with the highest levels detected in the *frd3-7 zif1* double mutants (**Figure 3D**). This indicates a more strongly induced constitutive Fe deficiency response in the double mutants, as confirmed by western blot quantification (**Figure 3D**). Under Zn excess, IRT1 abundance appeared slightly elevated in *frd3-7* and *frd3-7 zif1* (**Figure 3C**), but no statistically significant differences were observed among genotypes (**Figure 3D**).

**Figure 3.**
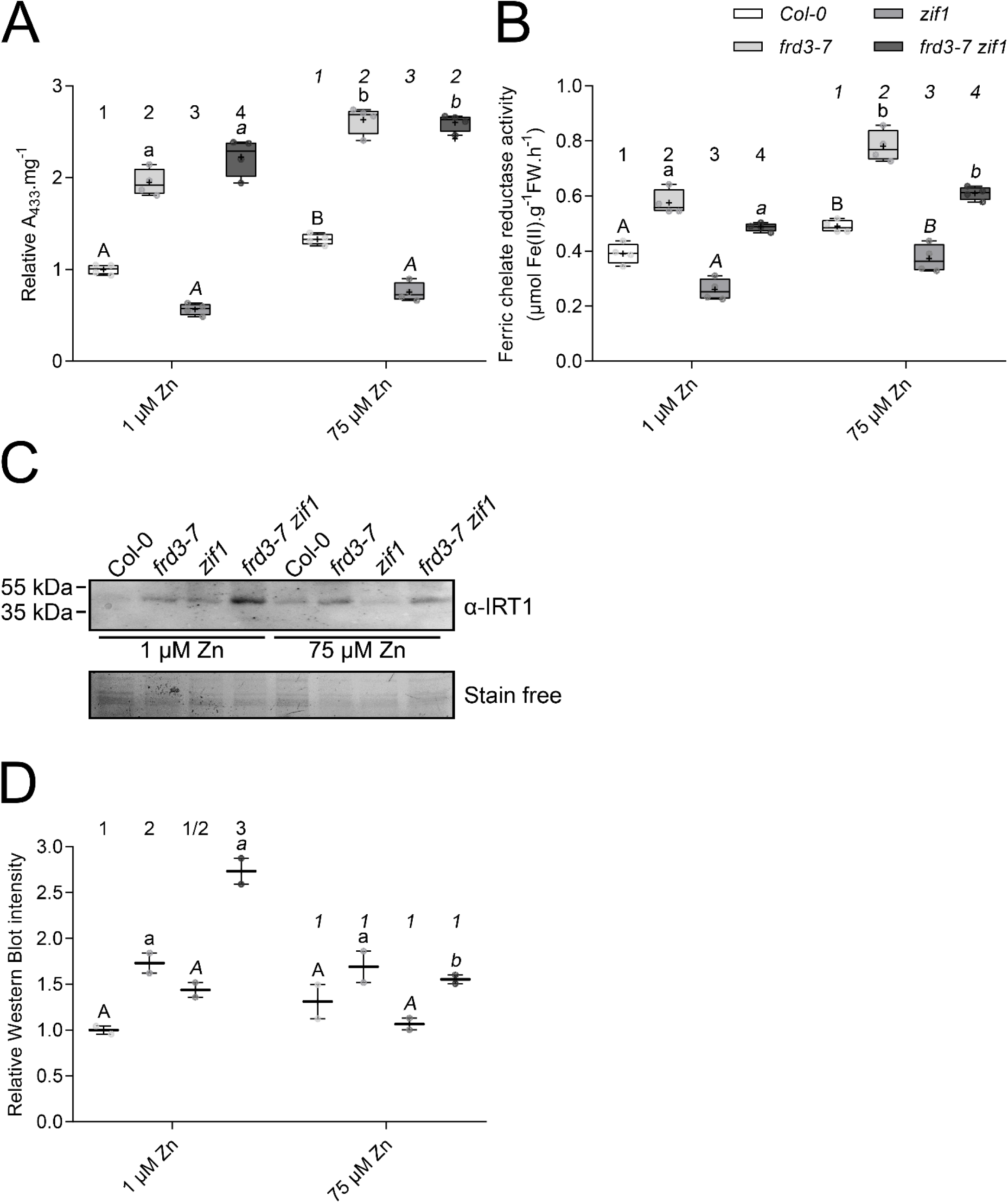
Fe uptake machinery activity in *frd3-7 zif1* plantlets under Zn excess. **(A)** pH assay and **(B)** Fe(III) chelate reductase activity in roots of Col-0 (white) *frd3-7* (light grey), *zif1* (middle grey) and *frd3-7 zif1* (dark grey) plantlets after 14 days of growth on solid media containing different Zn concentrations (1 and 75 µM ZnSO_4_). For pH assay, data are relative to Col-0 at 1 µM Zn. Box-and-whisker plots show the median (line), interquartile (box), and 1.5 interquartile (whiskers) of mean values from 4 independent experiments each including 3 series of 10 plantlets per genotype and condition. Mean values are represented by ‘+’ symbols. The data were analyzed by two-way ANOVA and followed by Bonferroni multiple comparison post-test. Statistically significant differences between conditions are indicated by letters (*P*<0.05) or between genotypes are indicated by digits (*P*<0.05). **(C–D)** Quantification of IRT1 protein abundance. **(C)** Representative western blot showing IRT1 protein accumulation in roots of Col-0, *frd3-7*, *zif1*, and *frd3-7 zif1* plantlets after 14 days of growth on solid media containing different Zn concentrations (1 and 75 µM ZnSO_4_). **(D)** Protein quantification of western blot analysis to monitor IRT1 protein levels in plantlet roots. Microsomal protein extracts were prepared from root tissues of Col-0 (white) *frd3-7* (light grey), *zif1* (middle grey) and *frd3-7 zif1* (dark grey) plantlets after 14 days of growth on solid media containing different Zn concentrations (1 and 75 µM ZnSO_4_). IRT1 was detected using anti-IRT1 antibodies (Agrisera). Stain-free imaging of total proteins served as loading control. Bars represent mean values from 2 independent biological replicates. The data were analyzed by two-way ANOVA and followed by Bonferroni multiple comparison post-test. Statistically significant differences between conditions are indicated by letters (*P*<0.05) or between genotypes are indicated by digits (*P*<0.05).

### The growth of frd3 zif1 double mutants is strongly impaired at older vegetative development stages

Having established how Zn excess and impaired metal chelation affect root system architecture, RAM integrity, and metal homeostasis in plantlets, we next examined vegetative growth at later stages to investigate how these early alterations influence growth, development and nutrient dynamics in older plants. Plants were grown for a total of six weeks under control (1 µM Zn) and zinc excess (20 µM Zn) conditions in hydroponics (**Figure 4A**). In control conditions, *frd3-7 zif1* double mutants showed severely reduced growth compared to the WT, with drastic decreases in root length, root biomass and shoot biomass (**Figure 4B-D and Supplementary Table S2)**. The *frd3-7* single mutant displayed a similarly strong growth defect, whereas *zif1* single mutants were largely comparable to the WT, except for a modest reduction in root length (**Figure 4 and Supplementary Table S2)**.

**Figure 4.**
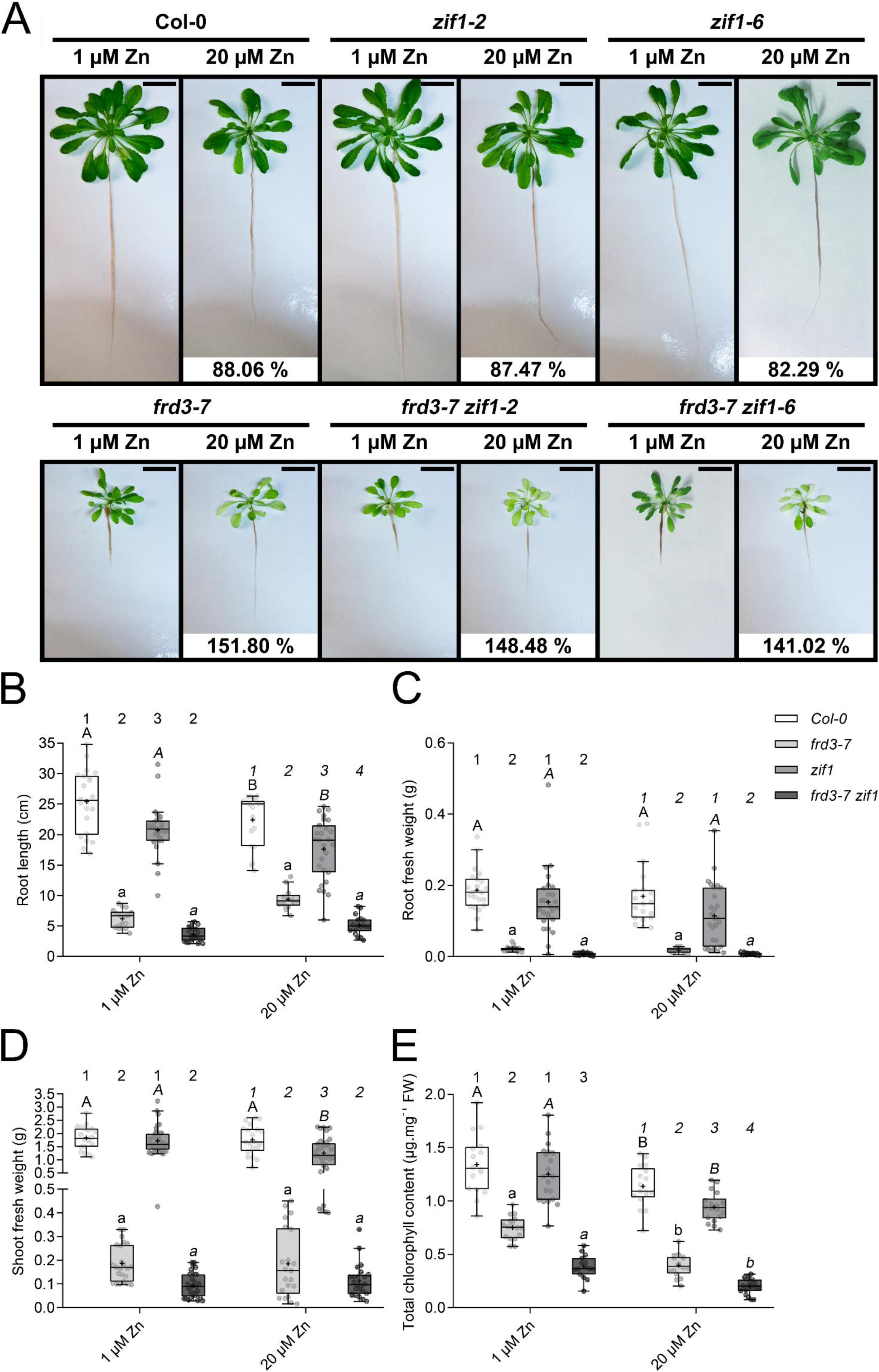
Zn excess partly rescues root growth of *frd3-7* and *frd3-7 zif1* mutants. **(A)** Col-0, *frd3-7*, *zif1-*2, *zif1-*6, *frd3-7 zif1-*2 and *frd3-7 zif1-*6 plant phenotypes upon growth in hydroponics and exposure to different Zn concentrations (1 and 20 µM ZnSO_4_). The numbers beneath the Zn excess (20 µM ZnSO_4_) panels for each genotype indicate the mean root growth ratio at 20 µM ZnSO_4_ relative to that at 1 µM ZnSO_4_, expressed as a percentage. The pictures are representative of 6 independent experiments. Scale bars: 2.5 cm. **(B)** Root growth, **(C)** root fresh weight per plant, **(D)** shoot fresh weight per plant and **(E)** total chlorophyll content per plant of Col-0 (white), *frd3-7* (light grey), *zif1* (middle grey) and *frd3-7 zif1* (dark grey) plants grown in hydroponics and exposed to different Zn concentrations (1 and 20 µM ZnSO_4_). Box-and-whisker plots show the median (line), interquartile (box), 1.5 interquartile (whiskers), and outliers (points beyond the whiskers) of values from 6 independent experiments each including 2-6 plants per genotype and condition. Mean values are represented by ‘+’ symbols. The data were analyzed by two-way analysis of variance (ANOVA) followed by Bonferroni multiple comparison post-test. Statistically significant differences between conditions are indicated by letters (*P*<0.05) or between genotypes are indicated by digits (*P*<0.05). All mean, median and relative values are provided in **Supplementary Table S2**.

Consistent with our previous report, Zn excess partially rescued the root growth in *frd3-7* (Scheepers et al., 2020), and a partial recovery was also observed in *frd3-7 zif1* double mutants, although growth remained strongly impaired relative to the WT at high Zn (**Figure 4A-B**). Conversely, Zn excess reduced root growth and biomass in the WT and in *zif1* single mutants (**Figure 4A-B**), with *zif1* showing decreased shoot biomass but no significant change in root biomass compared do the WT (**Figure 4C-D**).

The *frd3-7* and *frd3-7 zif1* all had reduced leaf chlorophyll contents in both growth conditions, but the *frd3-7 zif1* double mutants were the most strongly affected by high Zn (**Figure 4E**). This might be due to the cumulative defects of both *frd3* and *zif1* mutations, as *zif1* single mutants also appeared more chlorotic than the WT upon Zn excess (**Figure 4E**).

### High Zn exposure has distinct effect on metal distribution in frd3 zif1 double mutants

Similarly to the *frd3-7* mutant (Delhaize, 1996; Rogers and Guerinot, 2002; Durrett et al., 2007), the *frd3-7 zif1* double mutants had a very distinct ionomic profile compared to WT plants in control conditions, with Fe, Mn and Zn accumulation in root and shoot tissues (**Figure 5A-C and Supplementary Figure S5)**. In control conditions, *frd3-7* and *frd3-7 zif1* accumulated markedly higher levels of Fe (∼5.6- and ∼7.5-fold, respectively), Mn (∼15.5- and ∼13.9-fold, respectively), and Zn (∼5.6- and ∼10-fold, respectively) in roots compared with the WT (**Figure 5A-C and Supplementary Table S3)**. In contrast, metal accumulation in shoots increased only moderately in *frd3-7* and *frd3-7 zif1* **(Supplementary Figure S5 and Supplementary Table S3)**. Moreover, a visible Fe accumulation and H_2_O_2_ production in vascular tissues was observed in *frd3-7* and *frd3-7 zif1* (**Figure 5D**), which was in agreement with previous reports related to *frd3* mutants (Durrett et al., 2007; Scheepers et al., 2020; Fanara et al., 2022). In contrast, *zif1* single mutants presented a root ionomic profile, as well as Fe and H_2_O_2_ staining profiles comparable to WT plants (**Figure 5 and Supplementary Figure S5)**. At the shoot level, double mutants accumulated more Fe, Mn and Zn compared to the single *frd3-7* and *zif1* single mutants **(Supplementary Figure S5)**. These results were consistent with the accumulation of Fe and Mn in the *frd3-7 zif1* plantlets **(Supplementary Figure S3)**.

**Figure 5.**
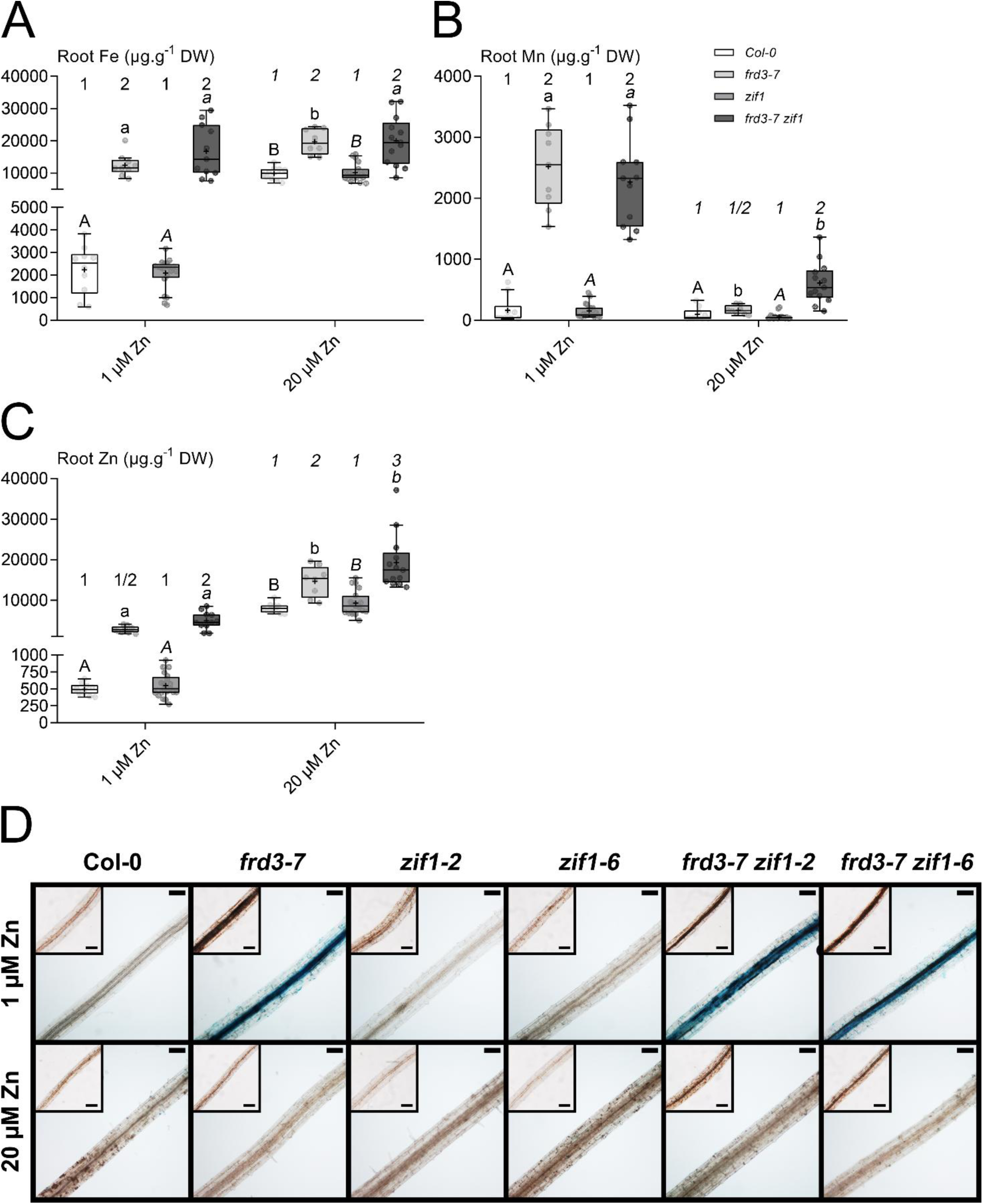
Metal accumulation and localization in roots of *frd3-7 zif1* double mutants. Metal accumulation [**(A)** Fe, **(B)** Mn and **(C)** Zn] in roots of Col-0 (white), *frd3-7* (light grey), *zif1* (middle grey) and *frd3-7 zif1* (dark grey) plants upon growth in hydroponics and exposed to different Zn concentrations (1 and 20 µM ZnSO_4_). Box-and-whisker plots show the median (line), interquartile (box), 1.5 interquartile (whiskers), and outliers (points beyond the whiskers) of values from 6 independent experiments, each including pools of 1-3 plants per condition and genotype. Mean values are represented by ‘+’ symbols. The data were analyzed by two-way ANOVA and followed by Bonferroni multiple comparison post-test. Statistically significant differences between conditions are indicated by letters (*P*<0.05) or between genotypes are indicated by digits (*P*<0.05). Shoot Fe, Mn and Zn accumulation data are shown in **Supplementary Figure S5**. DW, dry weight. All mean, median and relative values are provided in **Supplementary Table S3**. **(D)** Fe visualization by Perls’ staining (in blue) and H_2_O_2_ staining (in brown, insets) in root tissues of Col-0, *frd3-7*, *zif1* and *frd3-7 zif1* plants grown in hydroponics and exposed to different Zn concentrations (1 and 20 µM ZnSO_4_). Scale bars: 100 µm. The pictures are representative of 3 independent experiments.

High Zn exposure had distinct impact on the *frd3-7* single mutant and *frd3-7 zif1* double mutants. First, high Zn exposure only partially reduced Mn accumulation in roots of *frd3-7 zif1*, whereas it was fully reversed in *frd3-7* (**Figure 5B**). Second, the double mutant displayed higher Zn accumulation in roots at high Zn exposure (**Figure 5C**). Third, whereas Fe accumulation and H_2_O_2_ production in root vascular tissues were reversed by high Zn in the *frd3-7* mutant, as previously reported (Scheepers et al., 2020), oxidative stress was still observed in *frd3-7 zif1* independently of Fe accumulation (**Figure 5D**). This suggests that the H_2_O_2_ accumulation observed in roots of *frd3-7 zif1* upon Zn excess might likely be linked to the accumulation of other damaging metals (e.g., Mn and Zn) despite similar accumulation of Fe than what is observed in *frd3-7* (Remans et al., 2012; De Smet et al., 2015; Cuypers et al., 2016). In contrast, in the WT and *zif1* mutants, Zn excess had similar effect regarding root and shoot Fe, Mn and Zn concentrations, as well as Fe and H_2_O_2_ staining profiles (**Figure 5 and Supplementary Figure S5)**.

### High Zn exposure differentially modulates metal translocation in frd3 zif1 double mutants

Because both citrate and nicotianamine (NA) chelate metals and facilitate their intra-and intercellular mobility, as well as their long-distance transport, we investigated metal translocation. The *frd3-7* mutant accumulates citrate in root cells due to impaired xylem loading (Scheepers et al., 2020), whereas *zif1* mutants display elevated cytosolic NA levels as a result of defective vacuolar sequestration (Haydon et al., 2012). Metal translocation was therefore assessed by calculating shoot-to-root metal concentration ratios **(Supplementary Figure S6A-C)**.

In control conditions, *frd3-7* and *frd3-7 zif1* displayed strong reduction of shoot-to-root ratios for all three metals compared with the WT, indicating strong root retention and impaired translocation to shoots. This reduction was particularly pronounced for Fe and Mn, and was also evident for Zn. In contrast, *zif1* single mutants exhibited shoot-to-root ratios comparable to the WT for Fe and Zn, and only moderately reduced ratios for Mn **(Supplementary Figure S6A-C and Supplementary Table S3)**.

Upon Zn excess, shoot-to-root ratios decreased in the WT and *zif1* single mutants for Fe and Zn, reflecting a general reduction in translocation at high Zn. Nevertheless, *frd3-7* and *frd3-7 zif1* maintained lower Mn translocation compared with the WT, whereas *zif1* single mutants remained largely similar to the WT. Contrary to *frd3-7 zif1*, Mn translocation partially increased in the *frd3-7* single mutant under Zn excess compared with control conditions but remained substantially, yet not significantly lower than in the WT **(Supplementary Figure S6A-C and Supplementary Table S3)**.

Overall, these data indicate that altered NA compartmentation resulting from *ZIF1* disruption does not restore radial and long-distance metal partitioning in *frd3-7 zif1*. The persistently low shoot-to-root ratios in *frd3-7 zif1* demonstrate that NA redistribution is insufficient to compensate for defective citrate-dependent xylem loading, highlighting the non-redundant roles of these two chelation systems in root-to-shoot metal translocation.

### Defective NA compartmentation modulates the amplitude of Fe deficiency responses

Given the pronounced retention of Fe, Mn and Zn in roots of the *frd3-7* and *frd3-7 zif1*, we next questioned if the expression of genes known to substantially contribute to their transport into the root (e.g., *IRT1* and *NRAMP1*) was highly induced in these mutants. We therefore quantified transcript levels of selected Fe deficiency marker genes using quantitative reverse transcription polymerase chain reaction (qRT-PCR). We assessed the expression of genes encoding key components of the Fe uptake machinery (*AtAHA2*, and *AtFRO2*, and *AtIRT1*) (Martín-Barranco et al., 2020), their transcriptional activator *AtFIT* (Colangelo and Guerinot, 2004) (**Figure 6A-D**), and *AtNRAMP1* involved in Mn acquisition and contributing to Fe uptake (Castaings et al., 2016; Castaings et al., 2021) (**Figure 6E**). In WT and *zif1* single mutants, *AtFIT*, *AtAHA2*, *AtFRO2* and *AtIRT1* displayed low expression under control conditions and were induced upon Zn excess (**Figure 6A-D**). However, *zif1* single mutants exhibited a lower basal expression of *AtFIT* and *AtAHA2* and a marginaly lower expression of *AtFRO2* than the WT, and these differences persisted under high Zn. In contrast, *frd3-7* and *frd3-7 zif1* tended to display elevated expression of Fe uptake genes under control conditions, consistent with a constitutively activated Fe deficiency response as described in plantlets (**Figure 3**). Under high Zn conditions, *AtIRT1* expression reached levels comparable to those in the WT, whereas *AtFRO2* expression remained induced in *frd3-7* but was attenuated in the *frd3-7 zif1* double mutants. Although *AtAHA2* expression appeared induced in *frd3-7* and *frd3-7 zif1*, its expression level was not significantly different from that of the WT under Zn excess.

**Figure 6.**
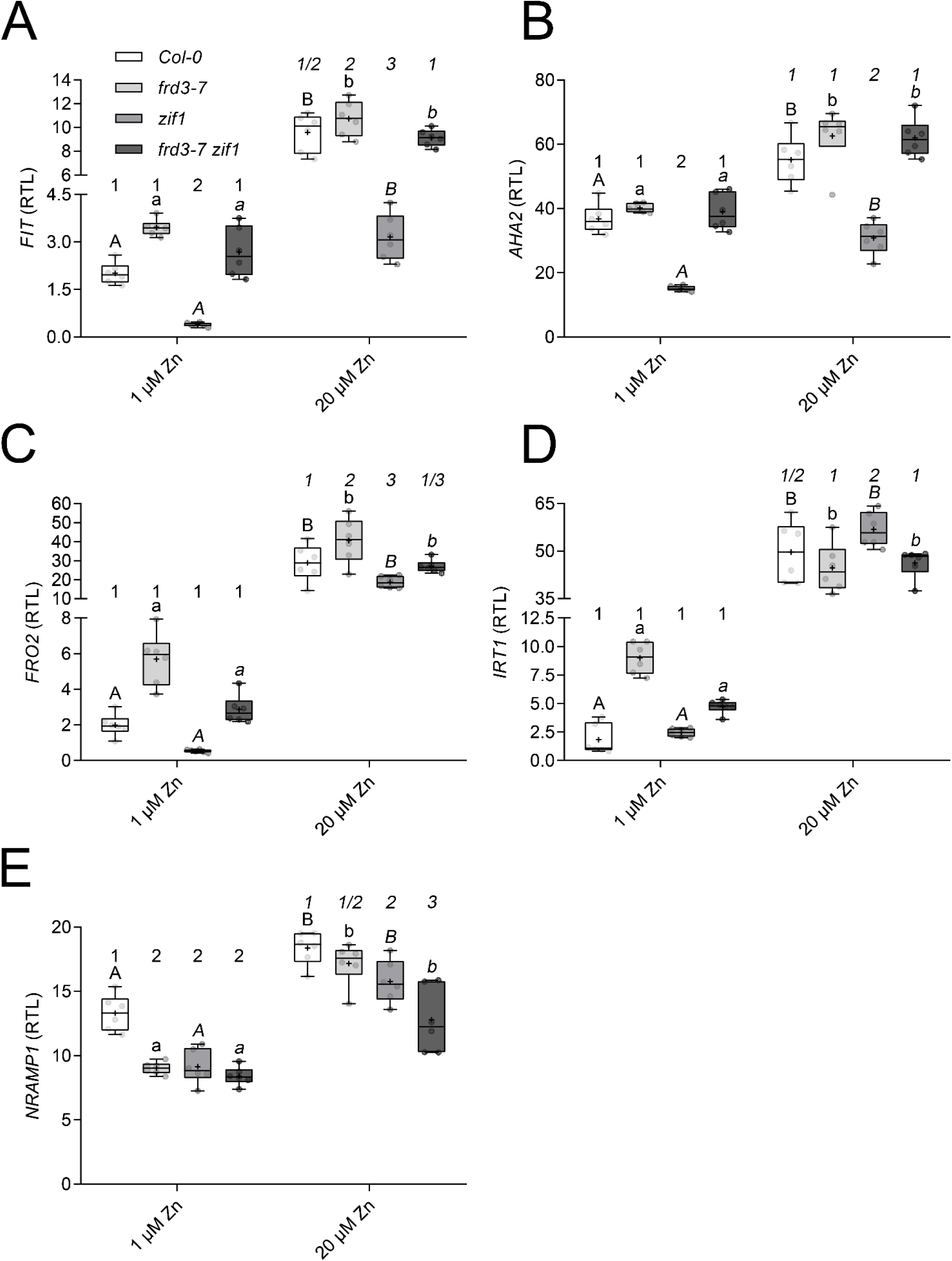
Expression analysis of Fe and Mn acquisition genes in *frd3-7 zif1* roots. Quantitative RT-PCR analysis of expression of the **(A)** *AtFIT,* **(B)** *AtAHA2,* **(C)** *AtFRO2,* **(D)** *AtIRT1*, and **(E)** *AtNRAMP1* genes in roots of Col-0 (white), *frd3-7* (light grey), *zif1* (middle grey) and *frd3-7 zif1* (dark grey) plants grown in hydroponics and exposed to different Zn concentrations (1 and 20 µM ZnSO_4_). Box-and-whisker plots show the median (line), interquartile (box), 1.5 interquartile (whiskers), and outliers (points beyond the whiskers) of values from 6 independent experiments, each including pools of 1-3 plants per condition and genotype. Mean values are represented by ‘+’ symbols. The data were analyzed by two-way ANOVA and followed by Bonferroni multiple comparison post-test. Statistically significant differences between conditions are indicated by letters (*P*<0.05) or between genotypes are indicated by digits (*P*<0.05). Values are relative to *AT1G58050*. RTL, relative transcript levels.

Finally, *AtNRAMP1* expression was reduced in all mutants under control conditions and remained significantly lower in *zif1* single mutants and *frd3-7 zif1* double mutants, but not in *frd3-7*, upon high Zn exposure (**Figure 6E**).

Together, these results indicate that the *frd3-7 zif1* largely retains the constitutive Fe deficiency signature of *frd3-7*, demonstrating that disruption of NA compartmentation does not override defective citrate-dependent signaling but instead modulates the amplitude of specific Fe uptake genes under Zn excess, fine-tuning the expression of selected components such as *FRO2*. In particular, *zif1* mutation leads to persistent downregulation of the Fe acquisition machinery-encoding genes, consistent with what was observed in plantlets (**Figure 3**), leading to intermediate transcriptional Fe deficiency responses displayed by *frd3-7 zif1* relative to each single mutant.

### NA biosynthetic genes are transcriptionally overexpressed in frd3 zif1 double mutants

As NA is a central chelator coordinating intracellular metal partitioning and long-distance transport, and as *nicotianamine synthase* (*NAS*) genes are tightly regulated by metal status (Klatte et al., 2009; Scheepers et al., 2020), we next investigated whether NA biosynthesis was transcriptionally altered in *frd3-7*, *zif1* and *frd3-7 zif1*.

Among the *AtNAS* genes, *NAS1* and *NAS2* were the most highly expressed in roots and were strongly overexpressed in *frd3-7* and *frd3-7 zif1* under control conditions, whereas their expression was lower in *zif1* single mutants **(Supplementary Figure S7A-B)**. By contrast, *NAS3* was the least expressed *NAS* gene and its transcript levels were slightly repressed by Zn excess in all genotypes **(Supplementary Figure S7C)**. *NAS4*, whose basal expression was intermediate relative to the other *AtNAS* genes **(Supplementary Figure S7D)**, was the most responsive to Zn excess, showing a ∼5.7-fold induction in WT roots **(Supplementary Figure S7E)**, consistent with a 48-hour exposure to Zn excess (Thiébaut et al., 2025b). While *NAS4* transcript levels were already elevated under control conditions in *frd3-7* and *frd3-7 zif1* relative to the WT, approximately twofold and fourfold inductions of *NAS4* in *frd3-7* and *frd3-7 zif1*, respectively, was observed relative to the WT in high Zn **(Supplementary Figure S7E)**. In contrast, *NAS2* showed the strongest overexpression under control conditions, with ∼7.3- and ∼4.7-fold higher transcript levels in *frd3-7* and *frd3-7 zif1*, respectively, compared with the WT. Upon high Zn exposure, this overexpression was attenuated in the *frd3-7* single mutant to levels similar to the WT, whereas it was further sustained in the double mutants **(Supplementary Figure S7B, E)**.

### Characterization of reproductive traits of frd3 zif1 double mutants

During prolonged hydroponic conditions, contrary to the other genotypes, the *frd3-7 zif1* double mutants were surprisingly not able to produce siliques **(Supplementary Figure S8A)**. The length of *frd3-7* or *zif1* siliques was marginally shorter than the one of the WT in control and Zn excess conditions **(Supplementary Figure S8A)**. Although the number of seeds per silique of *frd3-7* and *zif1* single mutants was comparable to the WT in control conditions, it tended to decrease upon Zn excess **(Supplementary Figure S8B)**. Seeds of *frd3-7 zif1* double mutants could only be obtained from plants grown on soil with a weekly sequestrene treatment. Because all seeds used in our assays originated from sequestrene-treated plants, we next analyzed the ionomic composition of these sets of seeds **(Supplementary Figure S8C-E)**. Fe levels were higher in seeds of *frd3-7 zif1* double mutants, a phenotype opposite to the lower Fe levels observed in seeds of each *frd3-7* and *zif1* single mutants **(Supplementary Figure S8C)**. In contrast, seed Mn levels were elevated in *zif1* single mutants **(Supplementary Figure S8D)**, while Zn levels were higher in all mutant seeds compared to the WT **(Supplementary Figure S8E)**.

## Discussion

Metal homeostasis in plants relies on a finely coordinated balance between uptake, chelation, intracellular compartmentation, intercellular mobility, and long-distance transport. Citrate and nicotianamine (NA) constitute two central chelators operating at distinct but interconnected levels: citrate predominantly support xylem loading and root-to-shoot translocation, whereas NA governs intracellular trafficking and remobilization from the phloem (Schuler et al., 2012). Genetic evidence from *nas4x*, and *frd3* mutants, as well as *ZIF1* overexpression lines indicates that each compound partially compensates for the absence of the other, but ultimately results in Fe deficiency symptoms, highlighting functional interdependence of these chelators (Klatte et al., 2009; Haydon et al., 2012; Schuler et al., 2012; Scheepers et al., 2020). Here, we combined impaired citrate xylem loading (*frd3-7*) with defective NA vacuolar sequestration (*zif1*) and characterized two independent *frd3-7 zif1* double mutants at different developmental stages, from vegetative to reproductive stages. Our study reveals that chelator compartmentation constitutes a core structural component of metal homeostasis robustness, linking ionomic changes with developmental alterations, including root system architecture and RAM integrity (**Figure 7**).

**Figure 7.**
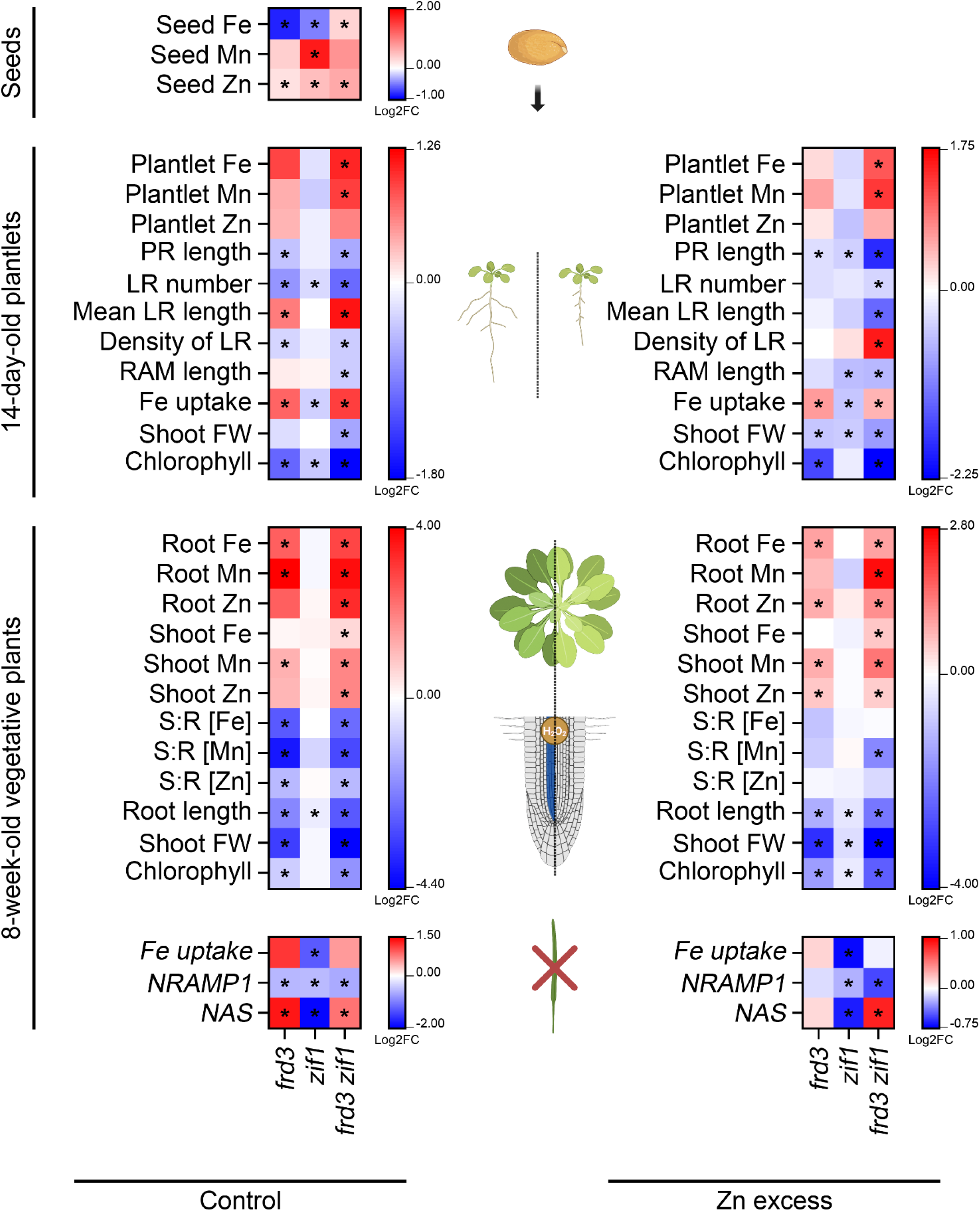
Integrated responses of the *frd3-7 zif1* double mutants to zinc excess compared with the wild-type. Under zinc (Zn) excess conditions, the root system architecture of the *frd3-7 zif1* double mutants was impacted compared with the wild-type: primary root length was reduced, with fewer and markedly shorter (1^st^ order) lateral roots, but greater density of lateral roots (**Figure 1**). The root apical meristem (RAM) size was reduced and contained fewer cortex cells, displaying visible cell death under severe Zn excess (**Figure 2**). Aerial tissues of the *frd3-7 zif1* double mutant plantlets developed less than the wild-type, with a drastic reduction of total chlorophyll content **(Supplementary Figure S2)**. Roots of the *frd3-7 zif1* double mutant plantlets showed higher activity of the Fe uptake machinery (AHA2, FRO2 and IRT1) than the wild-type (**Figure 3**), accompanied by an increased overall accumulation of iron (Fe) and manganese (Mn) **(Supplementary Figure S3)**. Under Zn excess in hydroponic conditions, mature aerial tissues of the *frd3-7 zif1* double mutants displayed more severe chlorosis symptoms than the wild-type (**Figure 4**), with increased levels of Fe, Mn and Zn **(Supplementary Figure S5)**. Roots of hydroponically grown *frd3-7 zif1* double mutants developed slightly better than in control conditions but remained shorter than the wild-type (**Figure 4**). They also contained higher levels of Fe, Mn and Zn, which was accompanied by visible oxidative stress (**Figure 5**). The translocation efficiency regarding Mn and Zn remained drastically impaired in the *frd3-7 zif1* double mutants **(Supplementary Figure S6)** despite a drastic increase in the expression of *nicotianamine synthase* (NAS) genes compared with the wild-type **(Supplementary Figure S7)**. The *frd3-7 zif1* double mutants also showed underexpression of the *NRAMP1* gene compared with the wild-type (**Figure 6**). The *frd3-7 zif1* double mutants were unable to produce any siliques under hydroponic conditions and could only develop them when grown on soil with a weekly treatment with sequestrene **(Supplementary Figure S8)**. Seeds obtained from these soil-grown, sequestrene-treated plants contained higher levels of Fe and Zn **(Supplementary Figure S8)**. Heatmaps summarize traits measured relative to the wild-type (log_2_ fold change, or Log2FC) in the corresponding conditions (control on the left, and Zn excess on the right). Color-coding indicates accentuated (red) or attenuated (blue) responses in the *frd3-7*, *zif1* and *frd3-7 zif1* mutants. Asterisks indicate statistically significant differences compared to the wild-type in the corresponding conditions. RAM length integrates both measurements expressed in micrometers and as the number of cortical cells in the cortex layer at 1 µM Zn and 150 µM Zn. The term “Fe uptake” represents the combined fold changes of AHA2, FRO2 and IRT1 activity and abundance in the plantlet-related panels, or the combined fold changes in the expression of *FIT*, *AHA2*, *FRO2* and *IRT1* genes in the hydroponics-related panels. Similarly, the term “NAS” represents the combined fold changes in the expression of *NAS1-4* genes. Plantlet and plant schemes illustrate the phenotypes of the *frd3-7 zif1* double mutants at both the shoot and root levels, including H_2_O_2_ accumulation and iron staining (blue in the vasculature). Fe: iron. Mn: manganese. Zinc: zinc. PR: primary root. LR: lateral roots. RAM: root apical meristem. FW: fresh weight. S:R: shoot-to-root metal concentration ([metal]) ratios. NAS: nicotianamine synthase. This figure was partly created with BioRender.com.

### Chelator imbalance compromises root system architecture and RAM integrity under Zn stress

Both nutrient deficiency and metal toxicity are known to reduce RAM size and promote differentiation (Richtmann et al., 2025; Thiébaut et al., 2025a; Thiébaut et al., 2025b). Zn levels in RAM are maintained across a wide range of external Zn supplies, and cell division persists without massive cell death, indicating the existence of protective allocation and controlled adaptation mechanisms safeguarding meristematic activity (Thiébaut et al., 2025a). In contrast, under severe Zn excess or Cd exposure, RAM shrinkage may involve oxidative stress and cell death. In *frd3 zif1* double mutants, Zn excess induces pronounced root system architecture (RSA) defects, reduced primary and lateral root growth (**Figure 1**), increased cell death in the RAM (**Figure 2**), and Zn hypersensitivity, all of which exceeded the developmental defects observed in either *frd3-7* or *zif1* single mutants. This reflects synergistic disruptions caused by impairments in both citrate and NA partitioning.

Although RAM length appears similar among genotypes, reduced cortex cell number and visible cell death suggest that cell cycle progression is altered and meristem function is compromised. Nutrient imbalance is known to indirectly modulate RSA through integrated transcriptional and developmental programs (Gruber et al., 2013; Giehl et al., 2014; van Dijk et al., 2022). Given the persistent H_2_O_2_ accumulation observed in *frd3 zif1* and the excessive Fe, Mn and Zn retention in roots (**Figure 5**), our results suggest that oxidative stress contributes to RSA defects likely induced by excessive root Mn and Zn levels (Remans et al., 2012; Zhao et al., 2017).

### Metal mis-partitioning triggers constitutive Fe deficiency responses despite Fe overaccumulation

A hallmark of *frd3 zif1* double mutants is the paradoxical coexistence of metal overaccumulation with strong activity of Fe uptake proteins, in particular AHA2 and IRT1, but in a lesser extent FRO2 (**Figure 3**). Similar transcriptional activation is observed under Zn excess (Thiébaut et al., 2025b), where Fe homeostasis genes are strongly induced despite no marked changes of Fe levels. This observation aligns with the idea that plants can respond to spatially or functionally partitioned metal pools, and not only to total tissue metal concentrations (Zlobin et al., 2019; Giehl et al., 2023; Lešková et al., 2025).

In *frd3 zif1* double mutants, impaired citrate export limits Fe delivery to shoots, likely triggering systemic Fe deficiency responses, similarly to what was observed in the *frd3* single mutant (Scheepers et al., 2020). Simultaneously, altered NA compartmentation disrupts intracellular metal distribution. This dual defect may generate local Fe misperception in specific root tissues, sustaining Fe deficiency responses (e.g., IRT1 accumulation) even under metal accumulation (**Figure 3 and Supplementary Figure S3)**. Moreover, the elevated Zn concentration in mutant seeds **(Supplementary Figure S8)** may contribute to the constitutive activation of Fe deficiency responses (**Figure 3**) upon radicle protrusion through competition with Fe uptake, as previously demonstrated for cadmium (Wu et al., 2012). Such constitutive Fe activation, including IRT1 which is thought to also substantially contribute to Zn uptake (Shanmugam et al., 2011; Merlot et al., 2021; Spielmann et al., 2022; Cointry et al., 2025), may further exacerbate Mn and Zn root uptake (**Figure 5**). Thus, chelator compartmentation appears critical not only for transport but also for accurate metal sensing and homeostatic responses.

### Hyperactivation of NA biosynthetic genes does not restore functional partitioning

The strong and sustained induction of *NAS* genes in *frd3 zif1* double mutant **(Supplementary Figure S7)** parallels observations under Fe deficiency or Zn excess, where *NAS* transcription responds to altered metal status (Klatte et al., 2009; Scheepers et al., 2020; Thiébaut et al., 2025b). However, increased NA biosynthesis does not restore radial or long-distance transport in the double mutant **(Figure S6 and Supplementary Table S3)**. This uncoupling indicates that NA abundance alone cannot compensate for defective compartmentation. Vacuolar sequestration via ZIF1 likely plays a regulatory role in modulating cytosolic NA pools and metal buffering capacity (Haydon et al., 2012). Indeed, *zif1* single mutants persistently inhibited the transcription of *NAS* genes compared to the WT, and the reduced relative *NAS* transcript levels under both control and Zn excess conditions likely aimed at limiting excessive cytosolic NA that might further perturbate metal-NA balance (such as Mn-NA chelates, (Pittman, 2005)). Although the vacuole represents the main intracellular Mn storage compartment and NA can chelate Mn to promote its mobility to the xylem vessels (Demirevska-Kepova et al., 2004; Pittman, 2005; Lei et al., 2007; Flis et al., 2016), putative elevated biosynthesis of NA combined to already elevated cytosolic NA levels in *frd3 zif1* double mutants did not efficiently restore metal root-to-shoot translocation **(Supplementary Figure S6)**. It is therefore evident that without proper compartmentation, elevated NA synthesis does not restore functional and spatial partitioning to sustain proper vegetative and reproductive development (**Figure 7**).

### Reproductive defects reveal systemic consequences of chelator miscoordination

The inability of *frd3 zif1* double mutants to produce siliques under hydroponic conditions underscores a potential systemic impact of disrupted chelator coordination. Seed ionome alterations further indicate that defective metal partitioning during vegetative growth propagates to reproductive tissues. Indeed, citrate primarily contributes to xylem-mediated translocation, whereas NA supports remobilization from phloem (Klatte et al., 2009; Schuler et al., 2012). The simultaneous disruption of both pathways likely compromised metal allocation during reproductive development, resulting in sterility. Thus, chelator compartmentation safeguards not only root growth but whole-plant reproductive capabilities.

### Developmental defects of frd3 zif1 demonstrate non-redundant and synergistic roles of citrate and nicotianamine chelators

The *frd3 zif1* double mutant reveals a synergistic, non-redundant, roles of citrate export into xylem vessels and NA compartmentation in maintaining metal homeostasis. Across developmental stages, *frd3 zif1* consistently exhibits accentuated phenotypes compared with either single mutant, including aggravated root system architecture defects, enhanced sensitivity to Zn excess, increased accumulation of Fe, Mn and Zn in whole-plantlets as well as in roots and/or shoots (especially under Zn excess), stronger IRT1 protein accumulation, persistent root oxidative stress, sustained *NAS* gene overexpression, and severe reproductive failure. Although some transcriptional responses to Fe deficiency in the double mutant appeared intermediate relative to the single mutants at older vegetative stages, metal distribution defects (including Fe and Mn accumulation in seeds) and developmental impairments are consistently more severe in *frd3 zif1*.

### Experimental procedures

#### Plant material and growth conditions

All experiments were conducted with *Arabidopsis thaliana* [accession Columbia-0 (Col-0)] as Wild-Type, *Arabidopsis thaliana frd3-7* mutant (Col-0 background) (Roschzttardtz et al., 2011), *zif1-2* [SALK_ 011408, (Haydon and Cobbett, 2007b)] and *zif1-6* mutant (SALK_058236, Col-0 background). *frd3-7 zif1* double mutants were obtained by crossing *frd3-7* and *zif1-2* or *zif1-6* homozygous plants. F1 lines were genotyped to check the presence of the T-DNA insertion in *ZIF1* in the *frd3-7* maternal background. F2 lines were also genotyped to identify double homozygous mutant lines. Two *frd3-7 zif1* lines, originating from each *zif1* parental line, were used in all experiments.

For hydroponics experiments, 2-week-old plantlets were transferred in hydroponic trays (Araponics, Liège, Belgium) and cultivated for 3 weeks in control Hoagland medium, followed by 3 weeks of experimental conditions. The nutrient medium was renewed with fresh solution once a week and 3 days before the harvesting. The control Hoagland medium included 10 μM FeIII-HBED [N,N′-di(2-hydroxybenzyl) ethylenediamine N,N′-diacetic acid monohydrochloride], 1 μM Zn (ZnSO_4_.H2O) and 5 μM Mn (MnSO_4_.H2O) as reported in (Talke et al., 2006; Hanikenne et al., 2008; Scheepers et al., 2020).

For plate experiments, surface-sterilized seeds were directly sown on square plastic Petri plates (Greiner Bio-One, Vilvoorde, Belgium) containing modified Hoagland supplemented with sucrose (1% w/v; Duchefa Biochimie, Haarlem, The Netherlands), ZnSO_4_ (1, 75 or 150 µM Zn), and agar (0.8% w/v, Agar Type M; Sigma-Aldrich). The plantlets were grown vertically after stratification.

All experiments were realized under 8 h light (100 µm photon m^-2^ s^-1^, 22°C)/16 h dark (20°C) in a climate-controlled growth chamber and were conducted at least thrice independently. Details on experimental replication are provided in figure legends.

#### Analysis of metal concentrations

Root and shoot tissues were harvested separately. Root tissues were desorbed and washed as described (Talke et al., 2006), and shoot tissues were rinsed in distilled water. Plantlets were desorbed and washed as described (Talke et al., 2006). After drying at 60°C for 3 days, 10-30mg of tissue were digested on a DigiPrep Graphite Block Digestion System (SCP Science) with 3mL HNO_3_ (≥65% v/v; Sigma-Aldrich) in DigiPrep tubes as described in (Scheepers et al, 2020). Element concentrations were determined by inductively coupled plasma-atomic emission spectroscopy (Vista AX, Varian).

#### Fe and hydrogen peroxide staining

For Fe Perls’ staining, root samples were vacuum infiltrated with HCl (4% v/v) and K-Ferrocyanide (II) (4% w/v; Sigma-Aldrich) at equal volume (1/1) for 10 min and incubated for 30 min at room temperature. The reaction was stopped by substituting the solution with distilled water (Roschzttardtz et al., 2009; Roschzttardtz et al., 2011). Images were taken using a binocular (SMZ1500; Nikon) equipped with a camera (Digital Sight DS-5M; Nikon).

Detection of H_2_O_2_ in root tissues was carried out according to (Baliardini et al., 2015). Briefly, root samples were vacuum infiltrated 15 min with 3-3’-diaminobenzidine tetrahydrochloride (1.25 mg/mL; DAB; Sigma-Aldrich), Tween-20 (0.05% v/v) and Na_2_HPO_4_ (200 mM) and incubated 45 min in the same solution. Then the roots were bleached in acetic acid/glycerol/ethanol (1/1/3) during 5 min at 100°C and then stored in glycerol/ethanol (1/4) before observation. Observation was performed using a binocular (SMZ1500; Nikon) equipped with a camera (Digital Sight DS-5M; Nikon).

#### Acidification capacity and ferric chelate reductase activity assays

Acidification capacity was assessed from a pool of five plantlets. Roots were incubated in 0.005% bromocresol purple (Roth, Karlsruhe, Germany) during 24 h in the dark, then the absorbance at 433 nm (A_433_) of the protonated form of the dye was then measured and expressed relative to the root weight of samples (Santi and Schmidt, 2009; El-Ashgar et al., 2012; Fanara et al., 2022).

The ferric (FeIII) chelate reductase activity was assessed from a pool of five plantlets. Samples were submerged in a reductase solution containing FeIII-EDTA (0.1 mM; Roth) and FerroZine (0.3 mM; Acros Organics, Morris Plains, NJ, USA) for 30 min in the dark. The absorbance at 562 nm (A_562_) of the FeII-FerroZine complex was then measured. The final calculation included the root weight of samples and the molar extinction coefficient of the complex (28.6 mm^-1^ cm^-1^) (Yi and Guerinot, 1996).

#### IRT1 Western blot

IRT1 western blots were performed using microsomal proteins fractions extracted from 200 mg of frozen plantlet root tissues as previously described (Martins et al., 2015; Spielmann et al., 2025). Microsomal proteins were separated on SDS-PAGE gel and transferred onto a PVDF membrane (ThermoScientific) using a methanol-based buffer. Proteins were detected using the polyclonal anti-IRT1 (Agrisera AS11 1780, 1:5000) and the HRP-conjugated anti-rabbit (Bio-Rad, 1:10 000) antibodies. Stain-free technology (Bio-Rad) was used as a loading control.

#### Chlorophyll content

Whole aerial parts from plantlets and hydroponically grown plants were weighted, then incubated in the dark during 72 hours or 7 days in ethanol (95%, VWR), respectively. Discolored leaves were removed, and the solution was then analyzed using spectrophotometer (Genesys20; Thermo Scientific), measuring absorbance at 649 nm and 665 nm. Total amount of chlorophyll were calculated using the following equation: Chl(*a+b*) = (6.1 A_665_ + 20.4 A_649_)/(fresh weight, in mg) (Wintermans and de Mots, 1965).

#### Automated acquisition of high-resolution pictures (Petrimaton)

The root system architecture of plantlets was assessed on high resolution pictures obtained using the Petrimaton apparatus (GDTech, Alleur, Liège, Belgium, and Quimesis, Wavre, Belgium) **(Supplementary Figure S9)**. The Petrimaton consists of an automated culture and light-treatment system for square Petri dishes (dimensions 120 x 120 x 17 mm). It includes: (1) a culture area accommodating up to 40 square Petri dishes equipped with dimmable (white) LED lighting; (2) a sealed area for individual and punctual light treatments equipped with Lumiatec PHS16 luminaries (GDTech, Alleur, Liège, Belgium) adjustable over 16 channels; (3) an infrared QR code detector; (4) a gripping and transfer system for square Petri dishes; and (4) an imaging area comprising a monochrome NIR line-scan camera with 8192 pixels per line (LA-GM-08K08A, Teledynedalsa, Waterloo, Canada) coupled to an APO-RODAGON-N 80/4.0 lens (Qioptiq, Goettingen, Germany) and an infrared backlight (peak at 850 nm). This device enables the implementation of potentially complex schemes of punctual light treatments and image acquisition for each individual Petri dish via QR code identification. As imaging is performed in the infrared range, the actinic effect on plant development is strongly limited.

#### RT-qPCR analysis

Upon harvesting, 14-day-old plantlets or plant root tissues were blotted dry, immediately frozen in liquid nitrogen and stored at -80°C. Total RNAs were prepared using 100 mg of homogenized tissues and NucleoSpin RNA Plant kit (Macherey Nagel, Düren, Germany) with on-column DNase treatment. cDNAs were synthesized with the RevertAid H Minus First Strand cDNA Synthesis Kit (Fisher Scientific) using Oligo(dT) and 1 µg of total RNAs. Quantitative PCR reactions were performed in a QuantStudio5 (Applied Biosystems, Waltham, MA, USA) using 384-well plates and Takyon Low Rox SYBR MasterMix dTTP Blue (Eurogentec, Seraing, Liège, Belgium) on material from at least three independent biological experiments, and a total of three technical repeats were run for each combination of cDNA and primer pair. Gene expression was normalized relative to *AT1G58050* as described (Pfaffl, 2001), as its expression was the more stable among all tested references (*AT1G58050*, *EF1α*, and *UBQ10*) (Czechowski et al., 2005). Primer reaction efficiencies were determined using LinRegPCR software (v2016.1) (Ruijter et al., 2009) **(Supplementary Table S4)**.

#### Mass spectrometry

Samples were prepared following previously published reports (Klatte et al., 2009; Haydon et al., 2012) and subsequently subjected to mass spectrometric analysis using a SYNAPT G2 HDMS mass spectrometer (Waters, Manchester, UK) as previously described (Scheepers et al., 2020). Frozen plantlet root tissues (60 mg) were ground to a fine powder with liquid nitrogen using a Retsch MM200 mixer mill (1 min at 25 Hz,). Metabolites were extracted by incubating the homogeneous powder in 500 µL of distilled water at 80°C, followed by agitation at 500 RPM for 15 min. This procedure was carried out twice to maximize recovery. Cell debris were finally eliminated by performing two successive centrifugation steps (20 min at 15,000 g each). Reference standards of citrate (1 mg mL^-1^, Sigma-Aldrich), malate (1 mg mL^-1^, Sigma-Aldrich) and nicotianamine (1 mg mL^-1^, Carbosynth) were prepared, and their mass-to-charge ratios (m/z) were determined prior to sample analysis to generate external calibration curves. Quantification of target metabolites in biological samples was achieved using the standard addition method. All analyses were performed on a SYNAPT G2 HDMS mass spectrometer equipped with an electrospray ionization source operating in negative ion mode. Tartaric acid (Sigma-Aldrich) was selected as an internal reference compound because it is not naturally present in Arabidopsis tissues (DeBolt et al., 2006; Scheepers et al., 2020). A stock solution was prepared in Milli-Q water (Millipore) at an exact concentration of 2.35 x 10^-4^ M. Using an analytical balance, 10 mg of tartaric acid and 100 mg of acetonitrile were added to 150 mg of each sample.

#### Statistical analysis

All data evaluation and statistics were done using GraphPad Prism 8 (GraphPad Software; version 8.0.1).

#### Accession numbers

The *Arabidopsis thaliana frd3-7* T-DNA insertion (SALK_122235), *zif1-2* T-DNA insertion (SALK_011408) and *zif1-6* T-DNA insertion (SALK_058236) lines were available in the Salk Collection.

## Author Contributions

MH: conceptualization and directing the research; SF and MSche: conducting most experiments with the help of MSchl and MSa for hydroponics and RT-qPCR, and from AF and PT for access to the Petrimaton apparatus; MB: performing mass spectrometry analyses; BB and MC: performing ICP-AES analyses; MH: supervision; SF and MSche: making the figures with the help of MB for mass spectrometry. SF, MSche and MH wrote the manuscript, with comments of all authors.

## Acknowledgements

We thank Prof. N. Verbruggen and Dr. J. Spielmann for helpful discussions. We thank Dr. C. Curie for the kind gift of *frd3-7* seeds. We thank A. Degueldre for his help with ICP-AES analyses. We acknowledge the use of BioRender.com for Figure 7. The authors wish to thank the COST ACTION 19116 PLANTMETALS for efficient networking and discussion. Funding was provided by the “Fonds de la Recherche Scientifique-FNRS” (PDR-T0120.18, PDR-T.0104.22 and CDR J.0009.17). S.F. and M.Sche. were doctoral fellows (‘Fonds pour la formation à la Recherche dans l’Industrie et dans l’Agriculture’).

## Conflict of interest

No conflict of interest declared.

---

**Supplementary Figure S1. Phenotypes of *zif1* single mutant plantlets exposed to Zn excess. (A)** Schematic representation of the *ZIF1* gene showing approximate location of 5’UTR and 3’UTR (light grey boxes), exons (dark grey boxes), and introns (black lines) constructed from information available at The Arabidopsis Information Resource (TAIR; www.arabidopsis.org). The location of the T-DNA insertions in the *zif1-2* and *zif1-6* lines are indicated. **(B)** Quantitative RT-PCR analysis of the expression of *AtZIF1* gene in roots of Col-0 (white), *zif1-2* (light grey) and *zif1-6* (dark grey) plants grown in hydroponics and exposed to control conditions (1 µM Zn). Data (relative transcript levels, RTL) represent mean values (±SEM) of 3 independent experiments, each including pools of 2 or 3 plants per condition and genotype each analyzed in 3 technical replicates. **(C)** Col-0, *zif1-*2 and *zif1-*6 plantlet phenotypes after 14 days of growth on solid media containing different Zn concentrations (1 and 75 µM ZnSO_4_). Scale bars: 1 cm. The pictures are representative of 6 independent experiments. **(D)** Primary root length, **(E)** shoot fresh weight per plant and **(F)** total chlorophyll content per plant of Col-0 (white), *zif1-2* (light grey) and *zif1-6* (dark grey) plantlets after 14 days of growth on solid media containing different Zn concentrations (1 and 75 µM ZnSO_4_). **(D)** Values are from 6 independent biological replicates each including 30-40 plantlets per condition. Middle line and whiskers represent mean values and standard deviations (SD), respectively. **(E-F)** Box-and-whisker plots show the median (line), interquartile (box), and 1.5 interquartile (whiskers) of mean values from 6 independent experiments each including 3 series of 10 plantlets per genotype and condition. Mean values are represented by ‘+’ symbols. **(D-F)** The data were analyzed by two-way analysis of variance (ANOVA) followed by Bonferroni multiple comparison post-test. Statistically significant differences between conditions are indicated by letters (*P*<0.05) or between genotypes are indicated by digits (*P*<0.05).

**Supplementary Figure S2. Shoot growth and chlorosis in *frd3-7 zif1* double mutants upon Zn excess. (A)** Shoot fresh weight per plant and **(B)** total chlorophyll content in shoots of Col-0 (white) *frd3-7* (light grey), *zif1* (middle grey) and *frd3-7 zif1* (dark grey) plantlets after 14 days of growth on solid media containing different Zn concentrations (1 and 75 µM ZnSO_4_). Box-and-whisker plots show the median (line), interquartile (box), and 1.5 interquartile (whiskers) of mean values from 4 independent experiments each including 3 series of 15 plantlets per genotype and condition. Mean values are represented by ‘+’ symbols. The data were analyzed by two-way ANOVA and followed by Bonferroni multiple comparison post-test. Statistically significant differences between conditions are indicated by letters (*P*<0.05) or between genotypes are indicated by digits (*P*<0.05).

**Supplementary Figure S3. Ionome of plantlets grown under Zn excess.** Metal accumulation [**(A)** Fe, **(B)** Mn and **(C)** Zn] in Col-0 (white), *frd3-7* (light grey), *zif1* (middle grey) and *frd3-7 zif1* (dark grey) whole plantlets after 14 days of growth on solid media containing different Zn concentrations (1 and 75 µM ZnSO_4_). Box-and-whisker plots show the median (line), interquartile (box), and 1.5 interquartile (whiskers) of values from 4 independent experiments each including 3 series of 8 plantlets per genotype and condition. Mean values are represented by ‘+’ symbols. The data were analyzed by two-way ANOVA and followed by Bonferroni multiple comparison post-test. Statistically significant differences between conditions are indicated by letters (*P*<0.05) or between genotypes are indicated by digits (*P*<0.05). DW, dry weight.

**Supplementary Figure S4. Metal chelator concentrations in root tissues**. **(A)** Citrate, **(B)** malate and **(C)** nicotianamine (NA) accumulation in Col-0 (white), *frd3-7* (light grey), *zif1* (middle grey) and *frd3-7 zif1* (dark grey) plantlets after 14 days of growth on solid media containing different Zn concentrations (1 and 150 µM ZnSO_4_). Box-and-whisker plots show the median (line), interquartile (box), 1.5 interquartile (whiskers), and outliers (points beyond the whiskers) of values from 3 independent experiments, each including pools of at least 48 plantlets per condition and genotype each analyzed in 3 technical replicates. Mean values are represented by ‘+’ symbols. The data were analyzed by two-way ANOVA and followed by Bonferroni multiple comparison post-test. Statistically significant differences between conditions are indicated by letters (*P*<0.05) or between genotypes are indicated by digits (*P*<0.05). Inset in **(C)**: the data were analyzed by one-way ANOVA (within conditions) and followed by Dunnett multiple comparison post-test

**Supplementary Figure S5. Metal accumulation in shoots upon Zn excess**. Metal accumulation [**(A)** Fe, **(B)** Mn and **(C)** Zn] in shoots of Col-0 (white), *frd3-7* (light grey), *zif1* (middle grey) and *frd3-7 zif1* (dark grey) plants grown in hydroponics and exposed to different Zn concentrations (1 and 20 µM ZnSO_4_). Box-and-whisker plots show the median (line), interquartile (box), 1.5 interquartile (whiskers), and outliers (points beyond the whiskers) of values from 6 independent experiments, each including pools of 1-3 plants per condition and genotype. Mean values are represented by ‘+’ symbols. The data were analyzed by two-way ANOVA and followed by Bonferroni multiple comparison post-test. Statistically significant differences between conditions are indicated by letters (*P*<0.05) or between genotypes are indicated by digits (*P*<0.05). Root Fe, Mn and Zn accumulation data are shown in **Figure 5**. DW, dry weight. All mean, median and relative values are provided in **Supplementary Table S3**.

**Supplementary Figure S6. Zn excess does not alleviate the reduction in metal translocation in *frd3-7* and *frd3-7 zif1* mutants.** Shoot-to-root metal concentration [**(A)** Fe, **(B)** Mn and **(C)** Zn] ratios from Col-0 (white), *frd3-7* (light grey), *zif1* (middle grey) and *frd3-7 zif1* (dark grey) plants grown in hydroponics and exposed to different Zn concentrations (1 and 20 µM ZnSO_4_). Box-and-whisker plots show the median (line), interquartile (box), 1.5 interquartile (whiskers), and outliers (points beyond the whiskers) of values from 6 independent experiments, each including pools of 1-3 plants per condition and genotype. Mean values are represented by ‘+’ symbols. The data were analyzed by two-way ANOVA and followed by Bonferroni multiple comparison post-test. Statistically significant differences between conditions are indicated by letters (*P*<0.05) or between genotypes are indicated by digits (*P*<0.05). All mean, median and relative values are provided in **Supplementary Table S3**.

**Supplementary Figure S7. *frd3-7 zif1* overexpresses *nicotianamine synthase* (*NAS*) upon both control and Zn excess conditions.** Quantitative RT-PCR analysis of the expression of **(A)** *NAS1*, **(B)** *NAS2*, **(C)** *NAS3*, and **(D)** *NAS4* genes in roots of Col-0 (white) *frd3-7* (light grey), *zif1* (middle grey) and *frd3-7 zif1* (dark grey) plants grown in hydroponics and exposed to different Zn concentrations (1 and 20 µM ZnSO_4_). Box-and-whisker plots show the median (line), interquartile (box), and 1.5 interquartile (whiskers) of values from 6 independent experiments, each including pools of 1-3 plants per condition and genotype. Mean values are represented by ‘+’ symbols. The data were analyzed by two-way ANOVA and followed by Bonferroni multiple comparison post-test. Statistically significant differences between conditions are indicated by letters (*P*<0.05) or between genotypes are indicated by digits (*P*<0.05). Values are relative to *AT1G58050*. RTL, relative transcript levels. **(E)** *NAS* genes expression relative to wild-type (Col-0) at 1 µM Zn, with numbers within each column representing fold-change in relative *NAS* expression.

**Supplementary Figure S8. Characterization of reproductive development traits of *frd3-7 zif1* double mutants. (A)** Silique length, and **(B)** number of seeds per silique from 12-week-old Col-0 (white), *frd3-7* (light grey), *zif1* (middle grey) and *frd3-7 zif1* (dark grey) plants grown in hydroponics and exposed to different Zn concentrations (1 and 20 µM ZnSO_4_). Box-and-whisker plots show the median (line), interquartile (box), and 1.5 interquartile (whiskers) of mean values from 4 independent experiments, each including pools of 1-3 plants per condition and genotype. Mean values are represented by ‘+’ symbols. The data were analyzed by two-way ANOVA and followed by Bonferroni multiple comparison post-test. Statistically significant differences between conditions are indicated by letters (*P*<0.05) or between genotypes are indicated by digits (*P*<0.05). n.a., not available as none of the *frd3-7 zif1* plants produced siliques under hydroponic growth conditions. Features were measured using the *straight-line* tool and the *cell counter* plugin on ImageJ/FIJI, respectively. **(C)** Seed Fe, **(D)** Mn and **(E)** Zn concentrations from Col-0 (white), *frd3-7* (light grey), *zif1* (middle grey) and *frd3-7 zif1* (dark grey) plants grown in soil and daily irrigated with water, as well as a weekly sequestrene supplementation (0.1 g L^−1^). Box-and-whisker plots show the median (line), interquartile (box), 1.5 interquartile (whiskers), and outliers (points beyond the whiskers) of values from seeds obtained from 8 to 27 individual plants. Mean values are represented by ‘+’ symbols. The data were analyzed by one-way ANOVA and followed by Tukey multiple comparison post-test. Statistically significant differences between genotypes are indicated by digits (*P*<0.05). DW, dry weight.

**Supplementary Figure S9. Pictures of the Petrimaton apparatus.** The root system architecture of plantlets (**Figure 1**) was assessed on high resolution pictures obtained using the Petrimaton apparatus (GDTech, Alleur, Liège, Belgium, and Quimesis, Wavre, Belgium) **(A)**. The Petrimaton consists of an automated culture and light-treatment system for square Petri dishes (dimensions 120 x 120 x 17 mm). It includes: **(B)** (1) a culture area accommodating up to 40 square Petri dishes equipped with dimmable (white) LED lighting; (2) a sealed area for individual and punctual light treatments equipped with Lumiatec PHS16 luminaries (GDTech, Alleur, Liège, Belgium) adjustable over 16 channels; (3) an infrared QR code detector; **(C)** (4) a gripping and transfer system for square Petri dishes; and (5) an imaging area comprising a monochrome NIR line-scan camera with 8192 pixels per line (LA-GM-08K08A, Teledynedalsa, Waterloo, Canada) coupled to an APO-RODAGON-N 80/4.0 lens (Qioptiq, Goettingen, Germany) and an infrared backlight (peak at 850 nm). This device enables the implementation of potentially complex schemes of punctual light treatments and image acquisition for each individual Petri dish via QR code identification. As imaging is performed in the infrared range, the actinic effect on plant development is strongly limited.

**Supplementary Table S1.** Mean, median and relative values of root system architecture traits in 14-day-old plantlets grown under control and Zn excess conditions.

**Supplementary Table S2.** Mean, median and relative values of root and shoot growth traits in hydroponically grown plants under control and Zn excess conditions.

**Supplementary Table S3.** Mean, median and relative values of root and shoot elemental concentrations under control and Zn excess conditions.

**Supplementary Table S4.** Quantitative RT-PCR primer sequences and primer efficiencies.

## Notes

### Competing Interest Statement

The authors have declared no competing interest.

