## Supplementary Figures S1-S9 for "Zinc and iron homeostatic interactions in a mutant lacking nicotianamine vacuolar storage and citrate xylem loading"

### Supplementary Figure S1

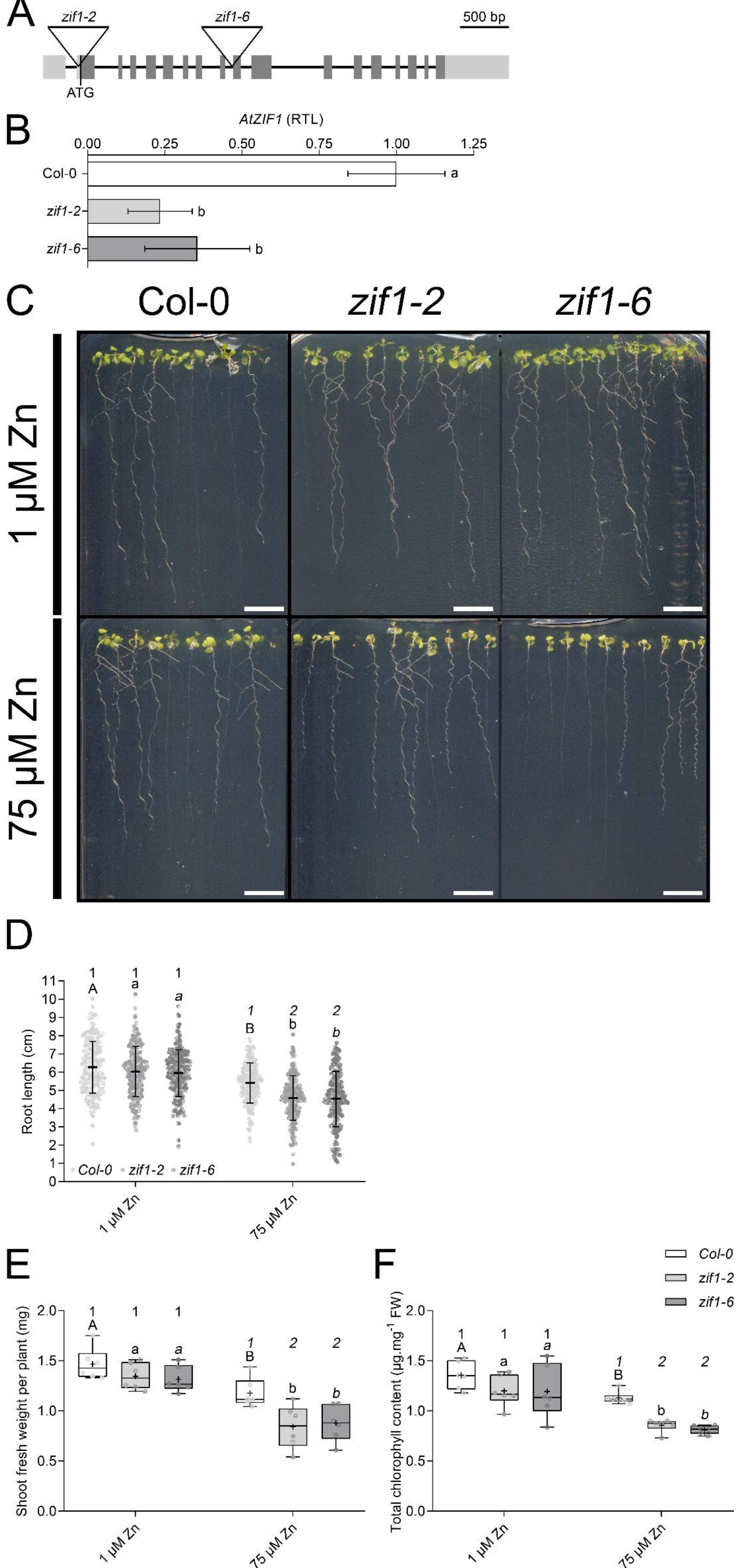

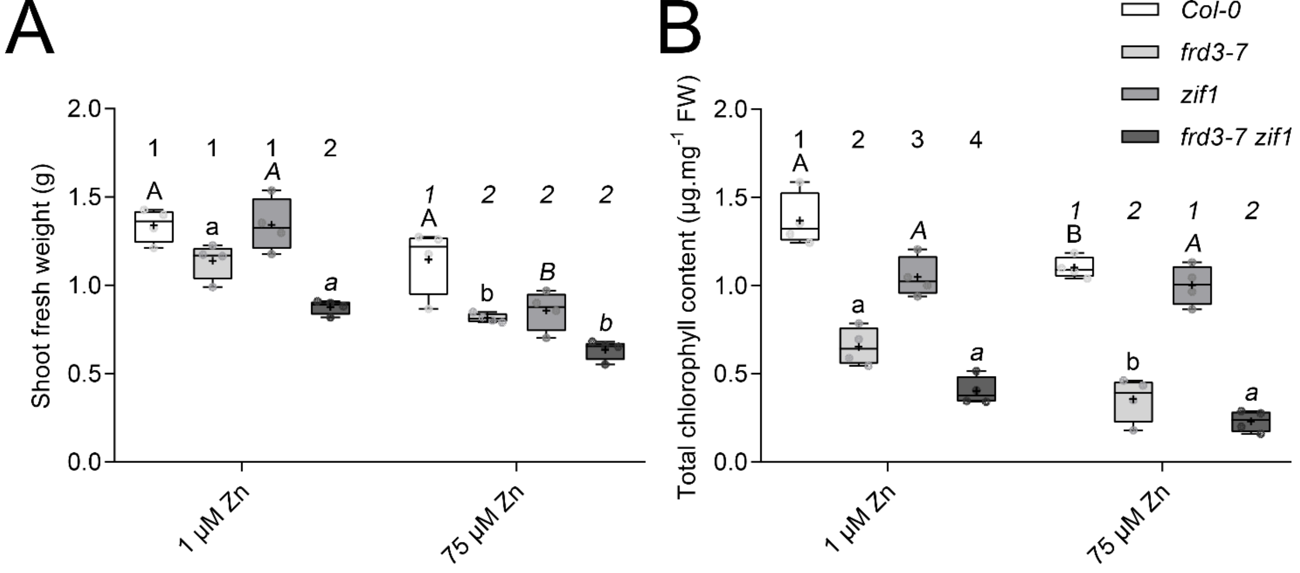


### Supplementary Figure S2

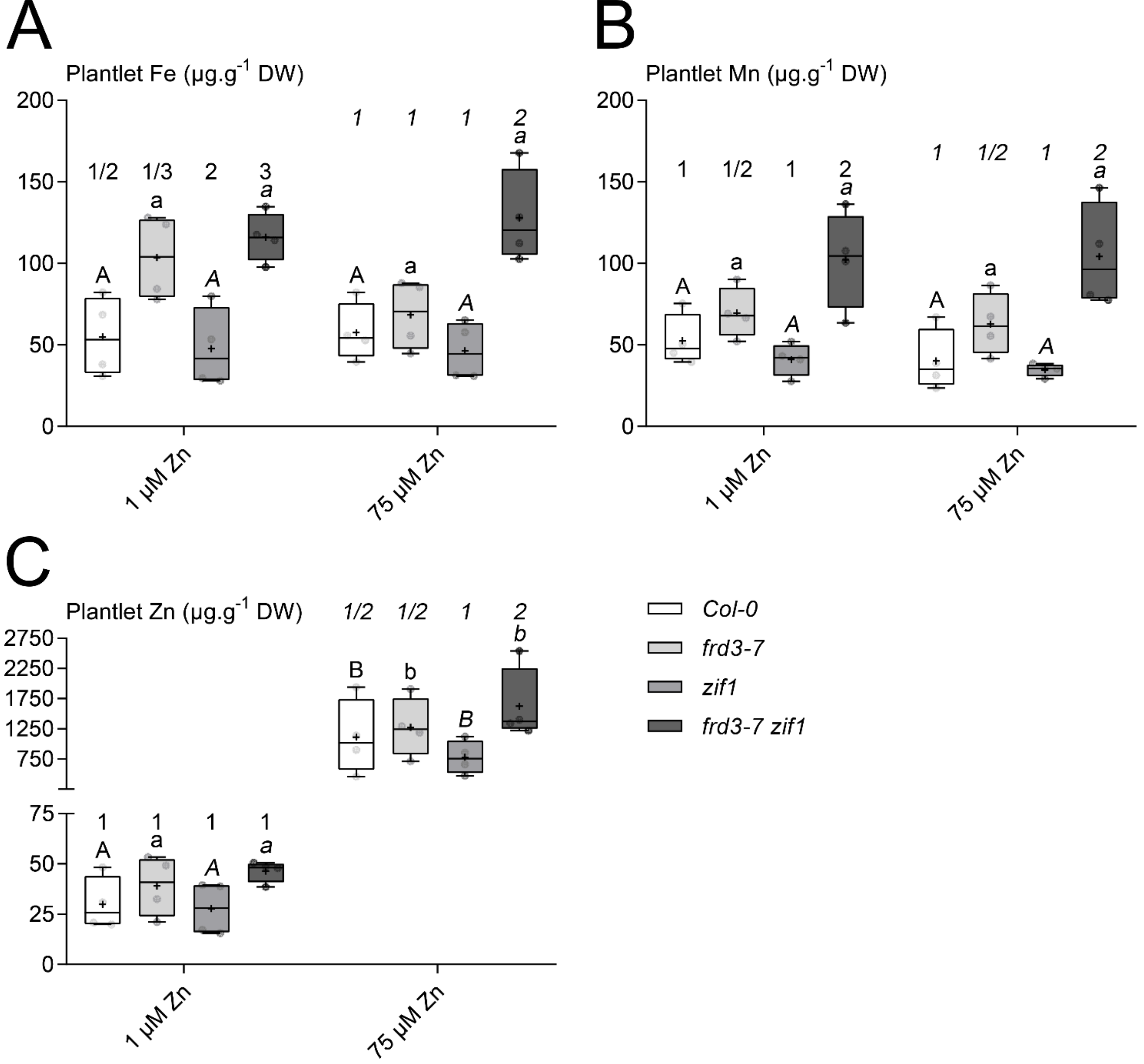


### Supplementary Figure S3

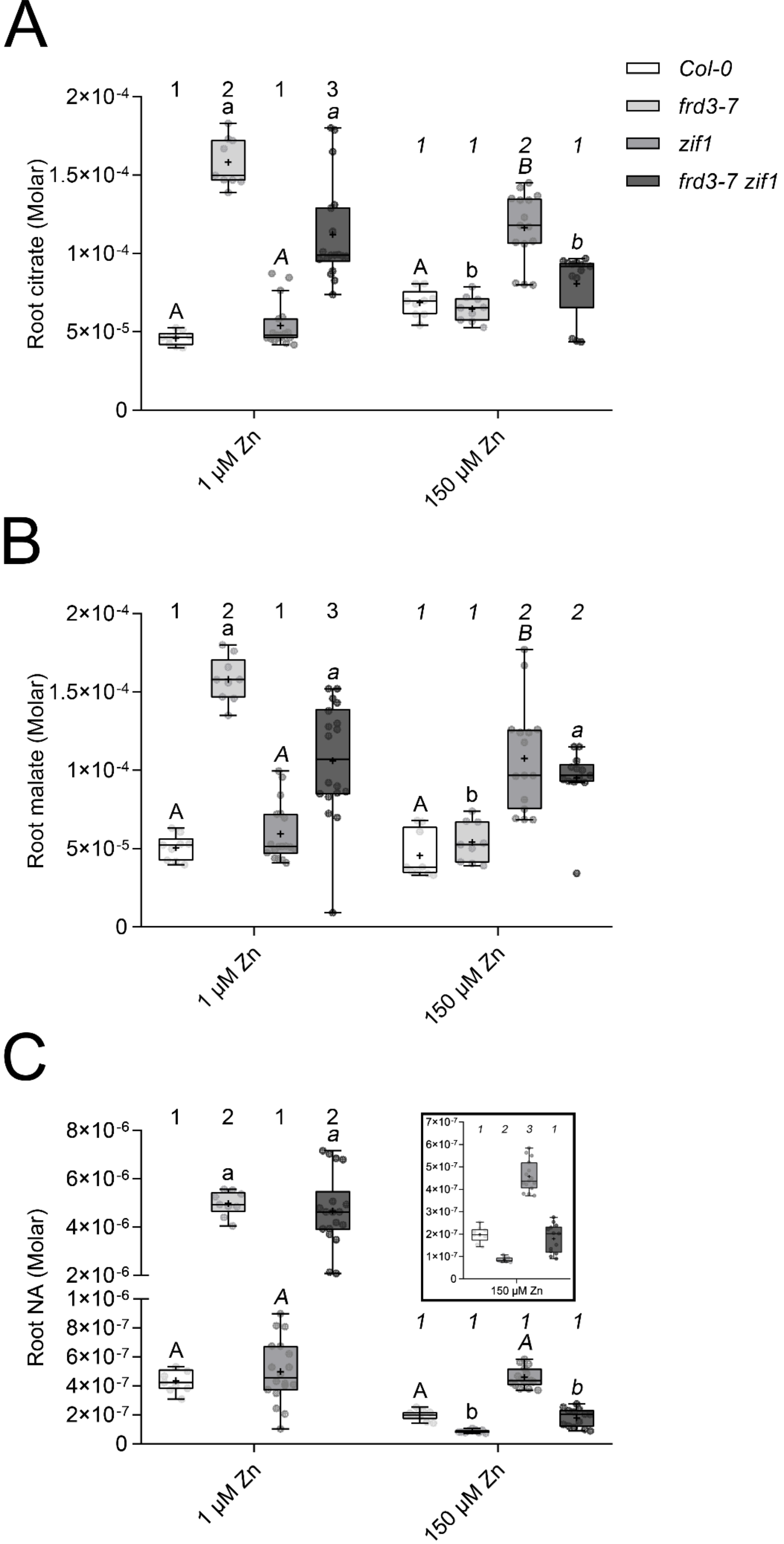


### Supplementary Figure S4

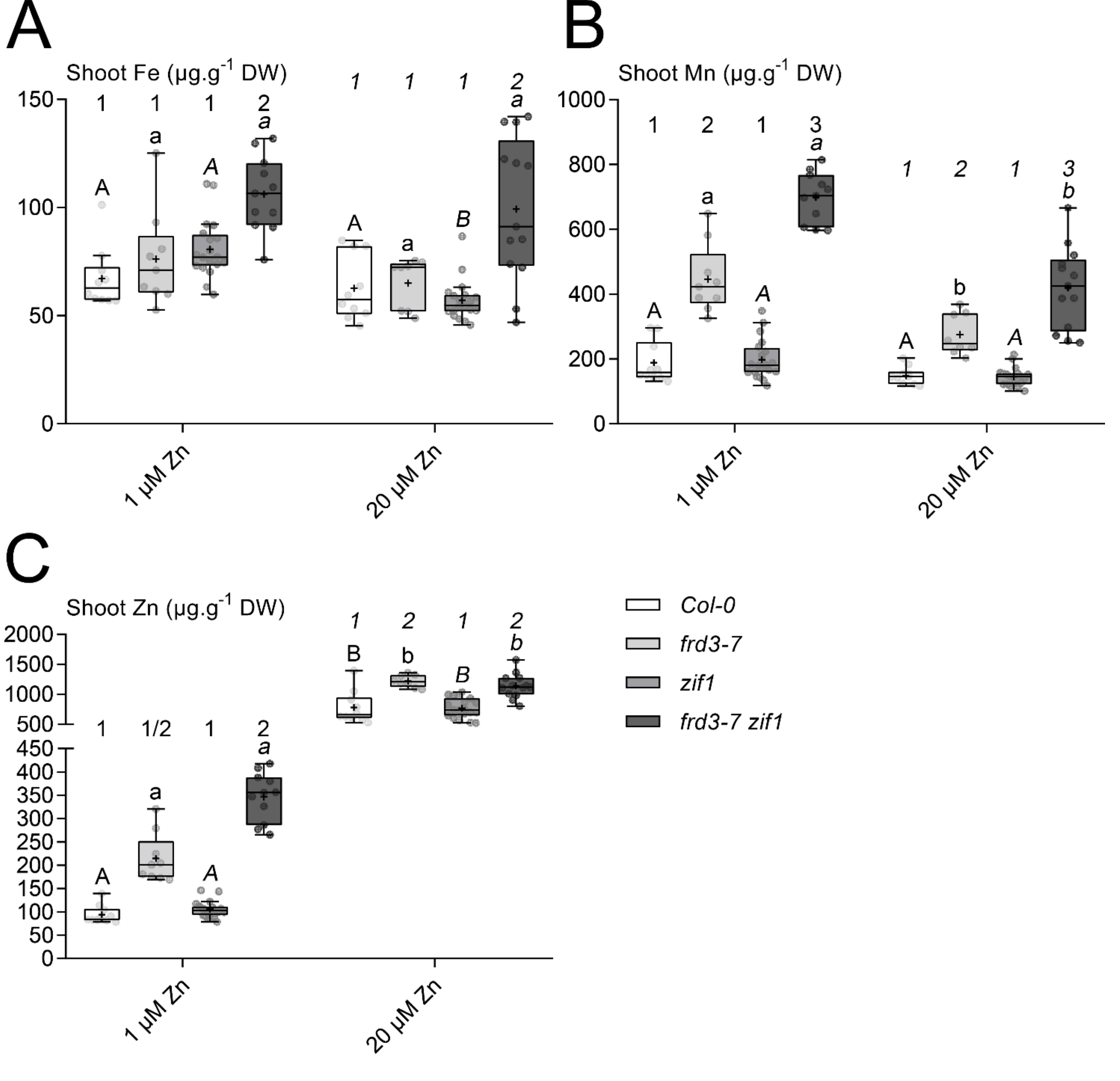


### Supplementary Figure S5

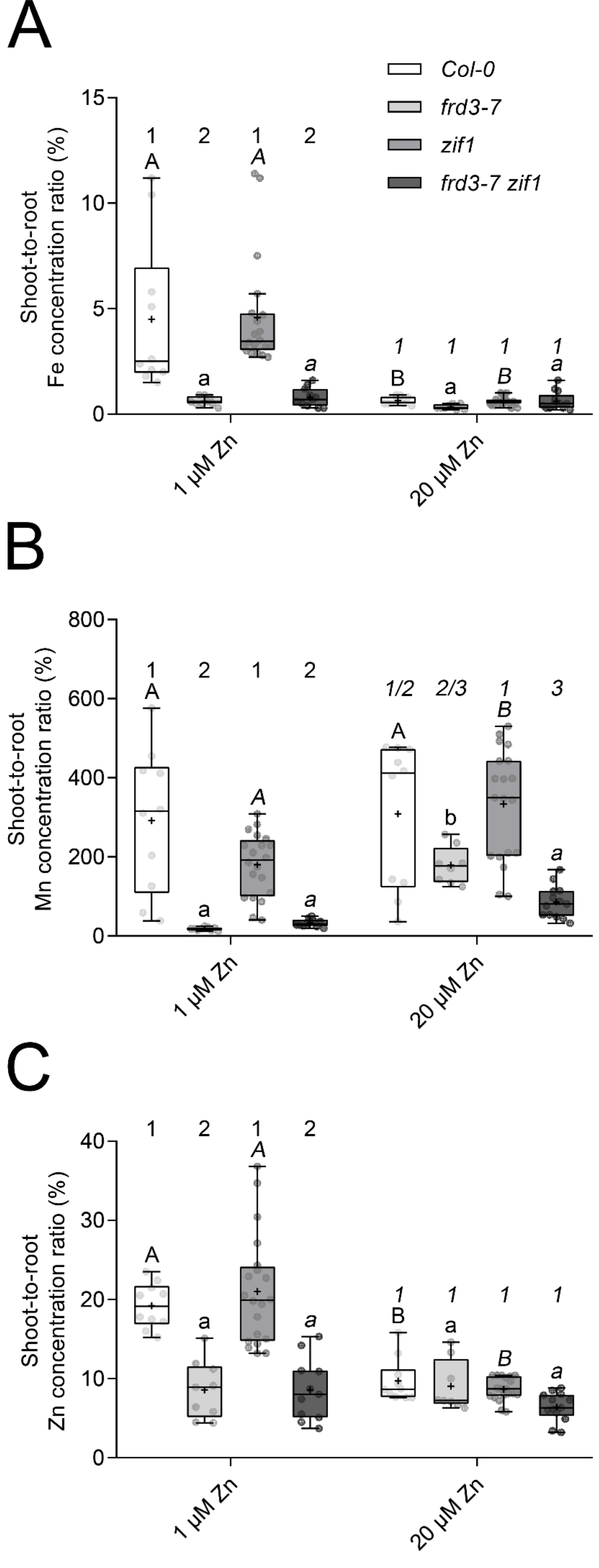


### Supplementary Figure S6

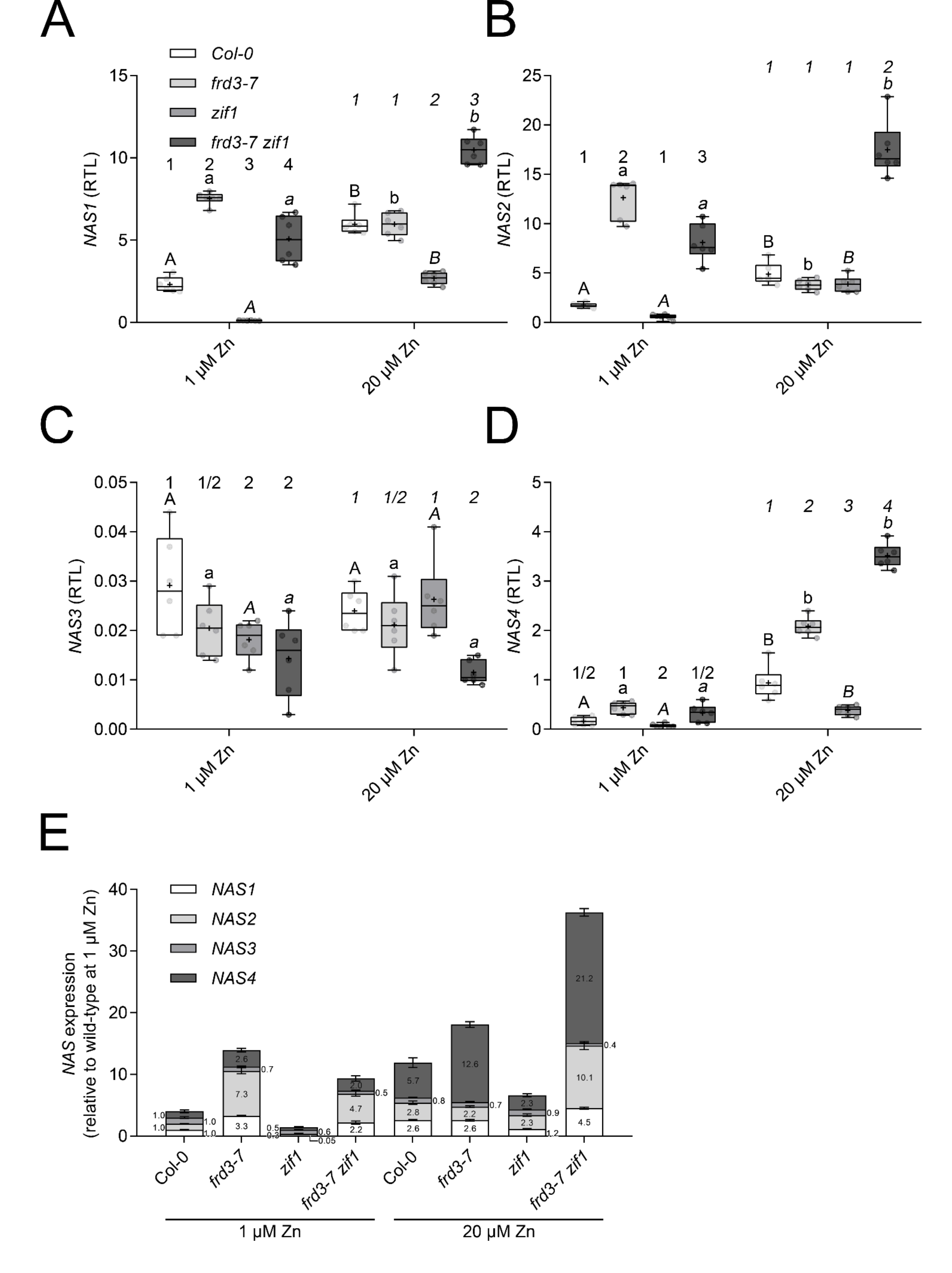


### Supplementary Figure S7

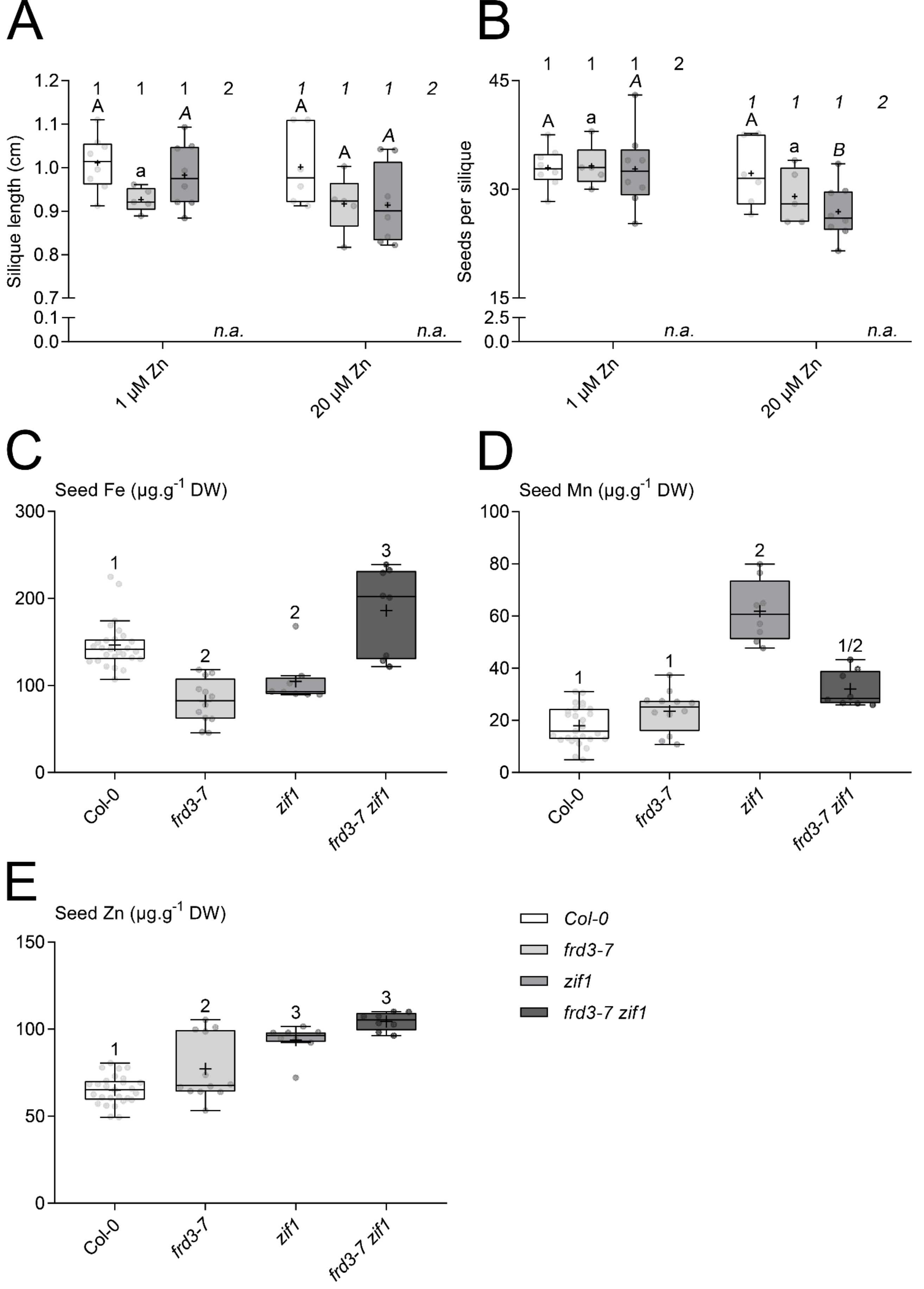


### Supplementary Figure S8

Supplementary Figure S9

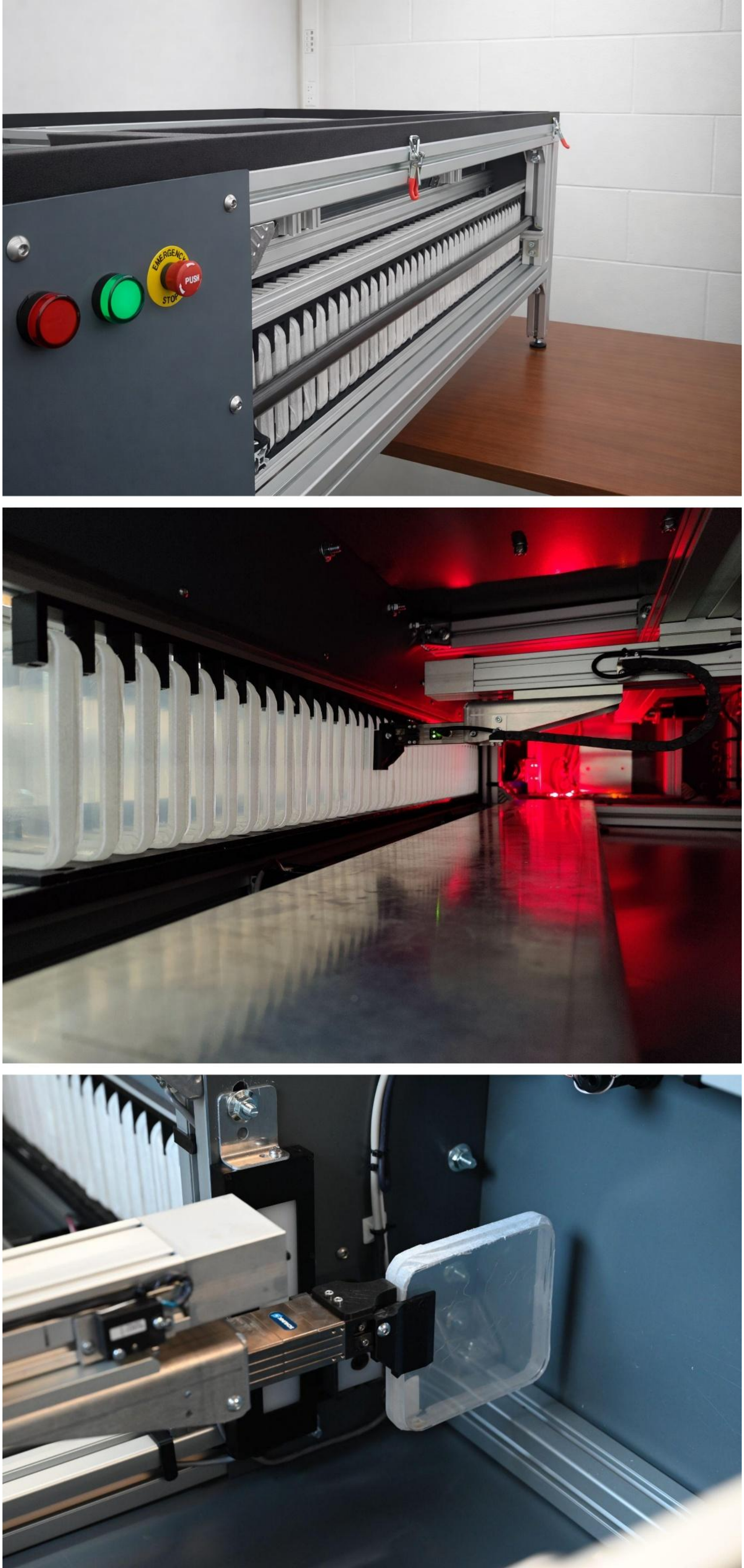


A

B

C
